# Emergent Statistical Laws in Single-Cell Transcriptomic Data

**DOI:** 10.1101/2021.06.16.448706

**Authors:** Silvia Lazzardi, Filippo Valle, Andrea Mazzolini, Antonio Scialdone, Michele Caselle, Matteo Osella

**Affiliations:** Physics Department, University of Turin and INFN, via P. Giuria 1, 10125 Turin, Italy; Laboratoire de physique de l’École normale supérieure (PSL University), CNRS, Sorbonne Université, and Université de Paris, 75005 Paris, France; Institute of Epigenetics and Stem Cells, Helmholtz Zentrum München; Feodor-Lynen-Strasse 21, 81377 München, Germany; Institute of Functional Epigenetics, Helmholtz Zentrum München; Ingolstädter Landstraße 1, 85764 Neuherberg, Germany; Institute of Computational Biology, Helmholtz Zentrum München; Ingolstädter Landstraße 1, 85764 Neuherberg, Germany

## Abstract

Large scale data on single-cell gene expression have the potential to unravel the specific transcriptional programs of different cell types. The structure of these expression datasets suggests a similarity with several other complex systems that can be analogously described through the statistics of their basic building blocks. Transcriptomes of single cells are collections of messenger RNA abundances transcribed from a common set of genes just as books are different collections of words from a shared vocabulary, genomes of different species are specific compositions of genes belonging to evolutionary families, and ecological niches can be described by their species abundances. Following this analogy, we identify several emergent statistical laws in single-cell transcriptomic data closely similar to regularities found in linguistics, ecology or genomics. A simple mathematical framework can be used to analyze the relations between different laws and the possible mechanisms behind their ubiquity. Importantly, treatable statistical models can be useful tools in transcriptomics to disentangle the actual biological variability from general statistical effects present in most component systems and from the consequences of the sampling process inherent to the experimental technique.

**Author summary:** Gene expression profiles represent how different cells use their genetic information. Similarly, books are specific collections of words chosen from a shared vocabulary, and many complex systems can be ultimately described by the statistics of their basic components. Leveraging on this analogy, we identified several emergent statistical laws in single-cell transcriptomic data that are universally found in complex component systems. A simple mathematical description sets these laws in a treatable quantitative framework and represents a useful tool for dissecting the different sources of gene expression variability.

## 1 Introduction

Almost every cell of an organism has the same gene content, but how these genes are expressed ultimately defines the cellular phenotype. If the gene repertoire is the genomic vocabulary, the transcription program represents how the different words are actually used by different cells to determine the cell identity [1]. Single-cell RNA sequencing (scRNAseq) technologies have recently given access to these cell-specific transcription programs[2], and large scale expression atlases have been compiled collecting thousands of single-cell expression profiles for all the major organs of different species [3, 4, 5].

The analogy between word statistics in a collection of texts and gene expression profiles in a large population of cells suggests that the transcriptome can be looked at as a complex component system [6]. Several complex systems of different nature and origin, from linguistics to biology, have an analogous modular structure with identifiable basic common building blocks that are used with different statistics. This statistics should contain information about the generative processes and the architectural constraints of the system. Books are composed by words, genomes of different species are collections of genes of different evolutionary families or associated to different biological functions, ecological niches are compositions of species with different abundances. Analogously, the transcriptional profiles of single cells are the sum of specific amounts of RNAs transcribed from a repertoire of common genes, and scRNAseq provides a picture of these profiles. The advantage of looking at single-cell transcriptomics as a complex component system is that a modelling framework and a set of analysis tools, based on statistical physics, have been developed for these systems. In fact, common statistical regularities have been characterized quantitatively in the different component systems described [7, 8, 9, 10, 11], and simple models have been proposed to explain their emergence. The first question we will address is if analogous statistical laws can be identified in large-scale transcriptomic data. While only few general regularities have been already recognised in transcriptomic data, this work presents the first detailed and systematic exploration across different datasets and experimental techniques. A quantitative systematic description is functional to the development and test of simple statistical models that can capture the connections between seemingly independent emerging laws. These data-driven models can in turn be used as “null models”, for example to disentangle genuine biological variability from technical or statistical effects in the context of transcriptomics. In fact, the observed variability in single-cell expression experiments typically has several possible sources, such as technical noise due to the experimental techniques, the intrinsic stochasticity in gene expression, and the biological variability actually setting the cell identity in terms of cell type and cell state (such as the cell cycle stage) [1, 12]. In particular, sampling noise associated with RNA capture and sequencing is inherent in RNA-sequencing techniques and can be a dominant source of noise, especially in single-cell transcriptomics where the starting RNA material is limited. This work focuses on cell atlases, as illustrative examples of large-scale scRNAseq datasets, with this complex systems perspective. We will identify emergent statistical laws in these datasets and assess their universality by comparison with properties of other component systems. Using a general and simple mathematical framework, we will show that several statistical properties of scRNAseq datasets can actually be explained as a result of the combination of heterogeneity in average expression levels and a sampling process. For example, we will show how this simple description can largely explain the empirical data sparsity in scRNAseq datasets, whose origin is still a debated topic in the field [13, 14], without invoking convoluted or ad-hoc assumptions.

On the other hand, some data features are not fully captured by this basic model, suggesting where to focus in order to extrapolate actual properties of biological variability. In fact, models based on empirical statistical laws, such as the one presented here or its potential future advancements, can be used as null models, for example to select the genes whose expression pattern is significantly divergent from the model expectation because of technical or biological reasons. We will show few illustrative examples of this model-driven gene-selection procedure. Analogously, the extant of the general discrepancy of a dataset properties from the model predictions should be related to intrinsic characteristics of the dataset, such as the degree of diversification of cell types. We will show that this indeed seems to be the case.

Finally, we will discuss how the addition of transcriptomic data to the increasing large set of systems displaying seemingly universal statistical properties is a relevant case study in the context of model generation. In complex systems theory, different general models and principles behind these ubiquitous laws have been proposed, and new empirical examples such as transcriptomic data, can be useful for model testing and selection. Therefore, this systematic exploration of statistical laws in single-cell transcriptomics could help to bridge the gap between mathematicians and physicists building general quantitative descriptions of complex systems, and computational biologists that could use the same descriptions to extract useful biological information from large-scale datasets.

## 2 Materials and Methods

### 2.1 Data sources

The Mouse Cell Atlas (MCA) was selected as the main illustrative dataset. In the MCA more than ~ 4 * 10^5^ single cells were profiled using scRNAseq from all major organs [3]. An advantage of this dataset is the use of Unique Molecular Identifiers (UMI) [15]. This technique allows the identification of the absolute number of unique RNA molecules detected by sequencing, thus eliminating the amplification noise. In the context of single-cell gene expression assays, this method provides a reliable estimate of the number of mRNA detected for coding genes and an estimate of the transcriptome size sampled. Our analysis therefore mainly focuses on absolute molecule counts, rather then relying on normalization techniques which are still a research area in sequencing data analysis.

We also analyzed the compendium of Tabula Muris (TM) for comparison. This atlas comprises an analogous number of cells from 20 organs and tissues [4] that were processed with the Smart-seq2 protocol [16], which produces a full-length transcriptome profiling but does not use UMI.

A dataset of bulk RNA-sequencing of healthy human tissues from the Genotype-Tissue Expression (GTEx) Project [17] was used to test the results on population-averaged transcriptomic data.

Finally, we analyzed two additional single-cell datasets, relative to a HEK cell line and to mouse fibroblasts, profiled with the recently introduced Smart-seq3 protocol [18].

### 2.2 The data structure for component systems

A transcriptomic dataset, and more generally a component system, can be described by a matrix 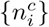 where each entry represents the counts relative to transcript (i.e, the component) *i* ∈ {1,…, *N*} in cell (i.e., the realization) *c* ∈{1,…, *R*}. *N* is the total number of different transcripts that could be present (the number of genes as a first approximation), which is essentially the vocabulary of our system. R is the number of cells analyzed. Each column of the data matrix is a vector 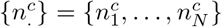 that fully describes the expression profile of a single cell c. The size of the transcriptome of a cell captured in the experiment is defined as 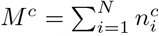. While in other component systems, such as texts of natural language, this parameter is simply the size of the realization (e.g., the book size), in our context *M^c^* represents the measured transcriptome size. Therefore, it does not necessarily correspond to the total number of transcripts in the cell because of the sampling process involved in RNA capture.

As described in the previous section, this work mainly focuses on two scRNAseq atlases: the MCA and the TM compendium. In these datasets, the total number of genes N (i.e., the genes with at least a single detected transcript) are respectively around 38 * 10^3^and 23 * 10^3^, while the number of cells R are 34 * 10^3^ and 41 * 10^3^. The distribution of transcriptome sizes *M^c^* is quite broad and dataset dependent. In the MCA, the average number of UMI per cell is ≃ 1 200 and the distribution is reported in S1.

### 2.3 An analytical framework to model gene expression data

This section develops the mathematical framework that will be used to describe the datasets. Using this framework, simple null models can be built to characterize the expected statistical behaviors given the model assumptions. The same mathematical description was applied in the context of metagenomic data [11], while an analogous approach has been previously introduced for scRNAseq data [19] and is at the basis of a recent Bayesian procedure for data normalization [20].

The key underlying assumption is that the observed mRNA counts 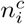 are the combined result of the inherent biological variability between cells and of the sampling process due to RNA capture and sequencing [11, 20]. Therefore, the probability of observing a specific expression profile 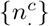 in a cell *c*, from which *M^c^* transcripts have been sequenced, is given by

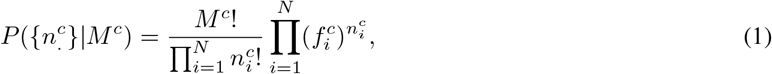

where 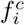 represents the true frequency of the mRNA *i* in cell *c*. The cell-to-cell variation in gene expression can be described by an unknown probability distribution *ρ*({*f*.}) setting the mRNA frequencies in the different cells. Thus, the probability of an expression profile in a cell with *M* observed transcripts becomes

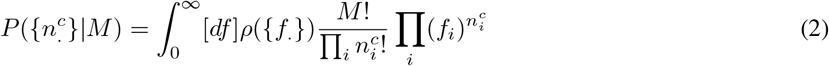

Focusing on a single gene *i* (by marginalizing the above expression), we have the probability of observing *n* counts as

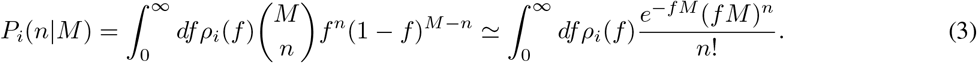

The Poisson approximation is valid when the number of mRNAs is large *M* ≫ 1 and even highly expressed genes occupy a small fraction of the total, which is typically the case in the datasets we want to analyze. The distribution *ρ_i_* captures the variability in expression of gene i due to both the different cell identities present in the cell population and the contribution from stochastic gene expression. On the other hand, the sampling variability is explicitly captured in the model by the binomial distribution. The average frequencies *f_i_* can be directly estimated by the empirical ones in the dataset [11]:

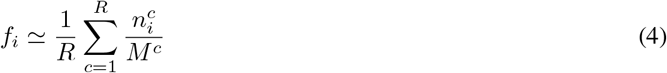

The ambitious goal would be to infer the distributions *ρ_i_*, and to distinguish the different contributions to the biological expression variability. Here, instead, we first consider a simple limiting case in which the actual biological variability is negligible with respect to the sampling noise. In this case, the distributions *ρ_i_* are extremely peaked with respect to the sampling noise and can be approximated with delta functions, i.e, *ρ_i_*(*f*) ≃ *δ*(*f* – *f_i_*). In this simplified scenario, the probability of observing an expression profile {*n*.}is given by the expression

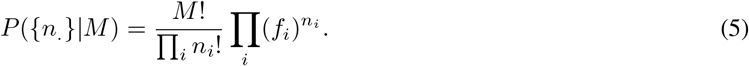

This model was previously analyzed to understand the origin of statistical regularities in different component systems [21].

We will compare the predictions of this simple model with empirical expression data. The idea is to understand what can actually be explained from the natural heterogeneity in average expression levels and from pure statistical effects due to the sampling process. One advantage of this model is that it is analytically treatable and provides mathematical predictions that can be directly tested against data. The situation of a dominant sampling noise can also be easily simulated. The transcript frequencies *f_i_* can be estimated from data with Eq. 4 and an ensemble of synthetic cells can be generated by randomly drawing *M^c^* transcripts, with *M^c^* values matching the empirical ones. The resulting surrogate datasets reproduce precisely the average expression levels and the sampling depth of the empirical dataset.

## 3 Results

### 3.1 Robust emergence of Zipf’s law for the gene expression levels at different scales

One of the hallmarks of complex systems, from real-world networks to natural language, is a high level of heterogeneity, which is often epitomized by the emergence of power-law distributions [22]. For component systems in particular, the frequency of components is often well described by a power law known as Zipf’s law [23, 22, 9, 6]. In natural language, this law describes the distribution of word frequencies in a corpus of texts, typically reported as a rank plot. In the context of transcriptomics, this would translate in a law for the distribution of gene expression levels in a large-scale dataset. Fig. 1A reports the rank-plot of the relative expression levels *f_i_* calculated by averaging across cells belonging to the same organ (different curves correspond to different organs) in the Mouse Cell Atlas (MCA) [3]. The distribution is largely compatible with a power-law decay with an exponent close to −1, as in the classic Zipf’s law, followed by an exponential tail. The shape of the distribution does not depend on the specific dataset or on the experimental technique used. An essentially identical plot (Fig. 1B) is obtained by looking at the same organs in an alternative mouse expression atlas, i.e., Tabula Muris [4], in which different sequencing methods were adopted. Even if the relative gene expression levels measured in the two atlases are correlated, the variability is substantial (S4). Besides biological variability, the two atlases adopt different protocols, therefore it is not so surprising that the measured expression levels are not perfectly conserved. However, Zipf’s law is robustly emerging. Also limiting the analysis to non-coding genes in the MCA, we still find the same distribution (S5).

**Figure 1:**
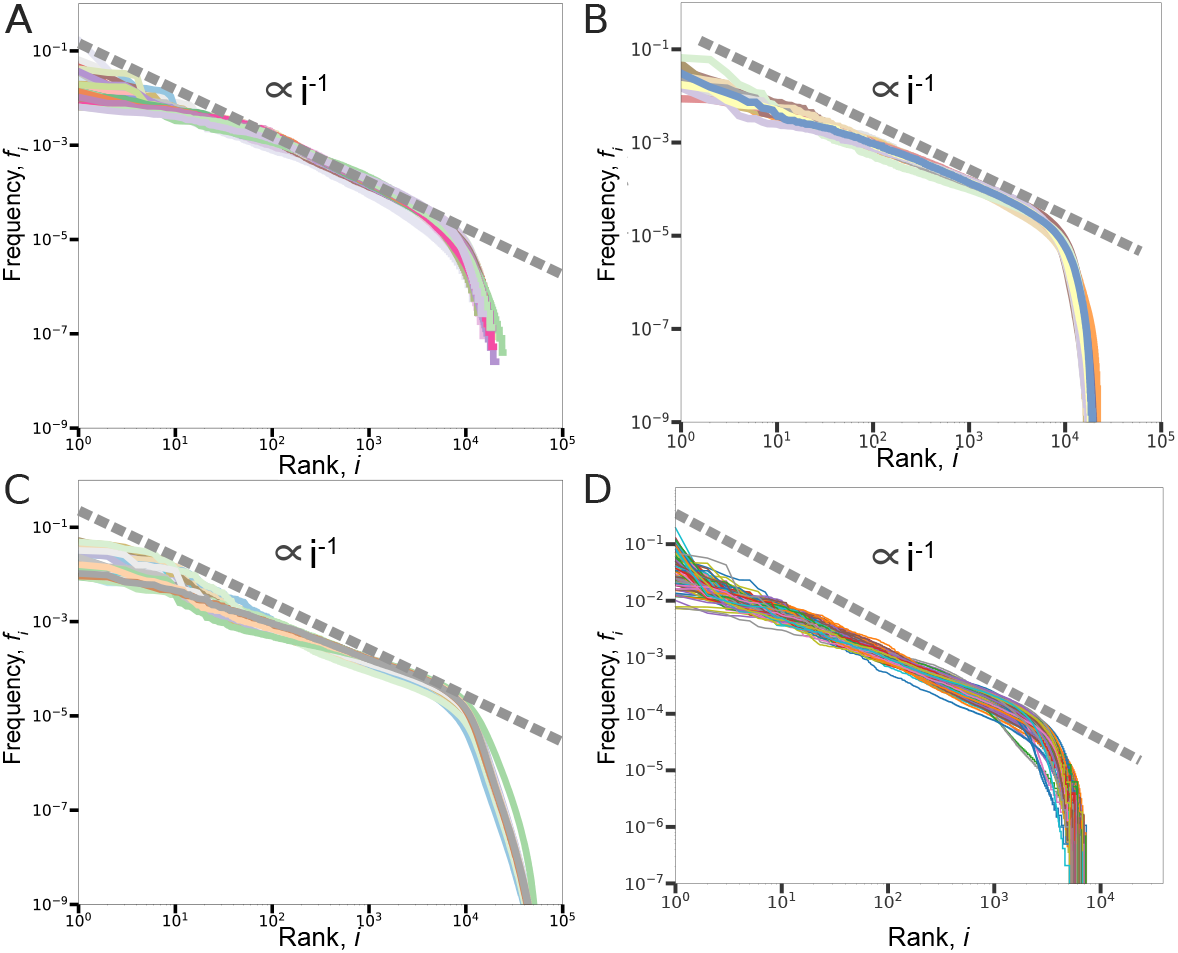
A robust Zipf-like law for gene expression levels. The average relative expression levels *f_i_* are estimated as described by Equation 4 and reported as a function of their rank. The distributions reported correspond to averages over single cells belonging to different mouse organs from the Mouse Cell Atlas **(A)**, from Tabula Muris database **(B)**, and to bulk RNA sequencing data from samples of healthy human organs in the GTEx database **(C)**. Each curve corresponds to a single organ or tissue and the corresponding color code is reported in S2.**(D)**The relative gene expression levels evaluated in single cells (without averaging) follow an analogous Zipf-like trend. We report the distribution relative to 100 cells from the heart sample in Tabula Muris. Similar results can be obtained from other organs or from the Mouse Cell Atlas (S3). The dashed lines are just a reference power-law scaling with exponent −1.

This statistical property seems indeed very general and not limited to scRNAseq data or to the specific species in analysis. For example, the same law emerges considering bulk RNA sequencing measurements across healthy tissues in human from the GTEx database [17] (Fig. 1C). This result corroborates previous observations based on microarray and SAGE (serial analysis of gene expression) datasets that reported a power-law distribution of gene expression levels across different species and experimental conditions [24, 25].

A natural question that can now be asked thanks to single-cell transcriptomics is if this emerging behaviour is a consequence of the averaging process or a property of the gene expression program of single cells. In other words, if gene expression in single cells displays actually a variety of cell-specific distributions that sum up in a Zipf-like law when averaged over a large cell population. Fig. 1D shows an illustrative example of the gene expression distributions in single cells. Besides some variability, the distributions recapitulate the population ones reported in the other panels. Therefore, the Zipf-like behaviour is an inherent property of single-cell expression profiles.

In conclusion, Zipf’s law appears to be a robustly emerging statistical property of gene expression data from bulk to single-cell experiments. This empirical law essentially sets the only free parameters *f_i_* of our null model described by Equation 5. Since this model only describes a sampling process given the empirical average frequencies of components, we can test what properties of the system can be explained merely by sampling effects and what features are instead potentially due to biological variability.

### 3.2 A Zipf’s law with multiple regimes

At a coarse grained view, the rank plot of gene expression levels can be described as a power law followed by an exponential tail. The presence of a double scaling in the component frequency distribution again is a general feature of several component systems. A similar behaviour can be observed by looking at protein domain frequencies in genomes of different species [6]. A double scaling was also observed in natural language [26], where it was tentatively explained by a model with two different classes of words: common words (high rank) composing a core vocabulary and the rest of more specific words in a vast vocabulary. Analogously, two different groups of genes can be distinguished in bulk transcriptomic data: a core of highly expressed genes with active promoters and a second group of lowly expressed and putatively nonfunctional transcripts [27]. This distinction was originally based on an observed bimodality in the histogram of expression levels in several bulk experiments. The same trend is present also in scRNAseq data from mouse organs (S6), and it is reflected in the drastic change of regime in the Zipf’s law, where the exponential tail contains the lowly expressed genes (Fig. 1). Fig. 2 shows that this exponential tail is also enriched in non-coding genes, that are indeed generally lowly expressed.

**Figure 2:**
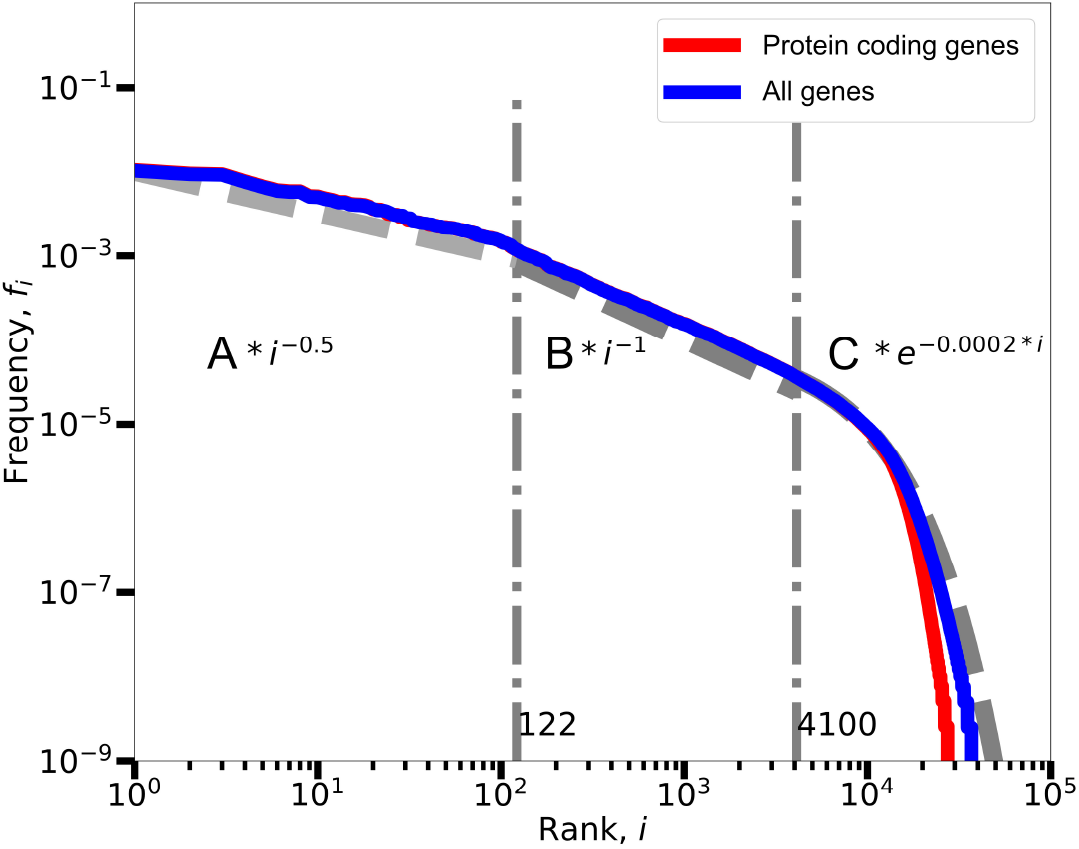
Multiple regimes in the rank-plot of the average expression levels. By considering the fitting error using a power law function in a window with variable width (S7), we were able to identify the part of the distribution well explained by a single power law, and consequently the two other regimes for highly expressed and lowly expressed genes. Excluding or including the non-coding genes from the analysis only influences the exponential tail of the distribution.

However, a more detailed and quantitative analysis indicates that also the top highly expressed genes deviate from the general power-law behaviour with exponent close to −1 (Fig. 2). In fact, it is possible to identify three different regimes that are approximately captured by two power laws with different exponents before the exponential tail. in order to quantitatively support these observations, we selected a window of ranks (e.g., identified by the dashed-dotted lines in Fig. 2) over which we performed a power-law fit of the frequencies. For different positions of the window boundaries, the coefficient of determination 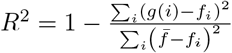 can be evaluated to select the range of ranks where the distribution is best explained by a power-law function. In the expression above, *g(i)* = *B* * *i*^−γ2^ is the power-law function of rank i obtained by fitting, while *f_i_* are the empirical frequencies with average 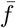 Once the boundaries and the best power-law fit of the central part of the distribution are defined by this procedure, the first regime (low ranks and high frequencies) can be fitted with an independent power-law function *g(i)* = *A* * *i*^−*γ*1^, and the third regime (high ranks and low frequencies) with an exponential function *g(i)* = *C* * *e*^−*γ3*i*^. S7 presents a more detailed illustration of this fitting procedure.

Considering all the cells in the MCA, highly expressed genes (around 100 genes) follow a power law with exponent close to −0.5, while the central part of the distribution is well described by an exponent close to −1 as in the classic Zipf’s law. Interestingly, a very similar law with three regimes was observed in a quantitative transcriptomic study of fission yeast [28]. The same behavior can be observed by considering the different tissues in the MCA separately (S8). Also the gene expression levels in single cells (Fig. 1D) seem to generally display three classes, but the higher level of fluctuations does not allow a refined analysis.

A Zipf’s law with three regimes emerges also across different datasets, as it can be qualitatively observed from Fig. 1. The precise values of the boundaries and of the fitted exponents are dataset-dependent as reported in the Supplementary S1 and S2. However, the general trend appear to be conserved: few highly expressed genes with a flatter expression distribution are followed by a central region of expression levels well described by a power law with exponent close to −1. Finally, the distribution shows an exponential tail for lowly expressed genes.

The highly expressed genes in the first regime belong to specific functional classes. For example, the most enriched Gene Ontology (GO) categories for the genes with rank lower than 100 in the MCA are associated to the basic protein translation processes (e.g, “structural constituent of ribosome” or “translation”) with Benjamini-corrected P-values lower than 10^−20^. GO enrichment analysis was performed using DAVID [29] and cross-checked using Metascape [30]. The lists of the most-enriched GO terms are reported in the Supplementary Materials (S3 and S4), where also the links to the full gene lists are presented. The genes in this first regime are quite common across organs, e.g., around 70%of them is in the top-100 highly expressed genes in at least half of the organs. In particular, 35 genes appear in the first regime of the 70%of the organs (S5). These genes present an enrichment for GO terms such as “ribosome” and “ribosome subunit” with P-values lower than 10^−20^. Therefore, the first regime appears to be composed by highly expressed genes related to basic functions that are common across different organs. This first core is followed by actively expressed genes that are more tissue specific and whose expression approximately follows the classic Zipf’s law with exponent −1.

### 3.3 The average number of detected transcripts follows Heaps’ law as predicted by a sampling process

A complex biological system such as an organ is composed by multiple cell types with transcription programs differentiated according to their functional role. Even the repertoire of genes that have to be transcribed is expected to vary from cell to cell as a function, for example, of the level of specialization of the cellular phenotype. Therefore, a basic observable difference between single-cell expression profiles could be the total number of genes that are actually transcribed. Resuming the analogy with texts of natural language, different texts typically use a different vocabulary (i.e., total number of different words), and the size of the vocabulary can depend on several factors such as the author style or the topic complexity. However, the average vocabulary of texts empirically displays a specific and well conserved sublinear scaling with the text size, known as Heaps’ law [31, 9, 32]. Again, an analogous law relates the number of different genes or protein domains to the genome size in prokaryotes [10]. Transcriptomic data present the additional complication that the number of detected transcripts also depends on the sampling process due to RNA capture. This naturally introduces a dependence on the sampling efficiency which is proportional to *M^c^*, i.e., the total number of captured transcripts from a cell c. Fig. 3A shows the number of different mRNAs as a function of the total number of UMI as an estimate of the total number of detected mRNAs. This analysis cannot be naturally applied in absence of UMI, since a reliable measure of the sample size is needed.

**Figure 3:**
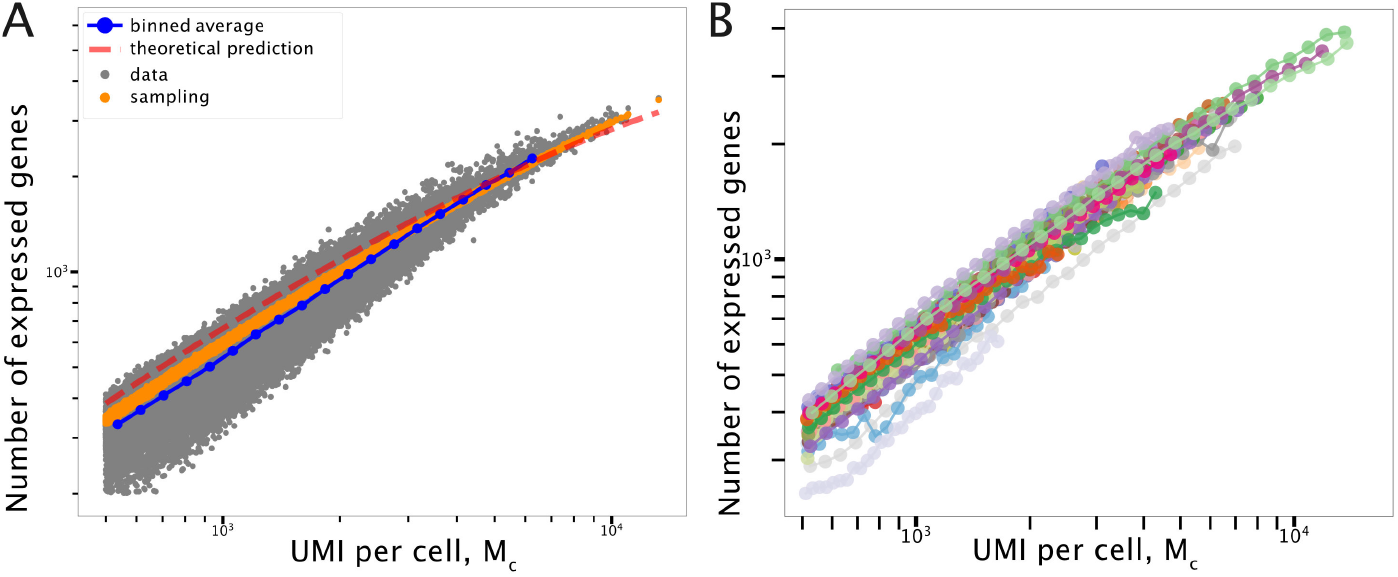
The number of detected different transcripts follows Heaps’ law. **(A)** The number of mRNAs with at least one detected transcript *h(M)* is reported as a function of the transcriptome size as measured by the number of UMI, i.e., 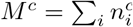. Each point in the scatter plot thus corresponds to a single cell, for the illustrative example of cells in the Bone Marrow from the MCA. The average empirical sublinear scaling (blue dots) is compared to the results of a stochastic sampling process using detailed simulations (orange dots) and analytical predictions (red dashed line). **(B)** The same sublinear average scaling is approximately conserved in all organs reported in the MCA.

The sublinear power-law scaling is very similar to the one found in other component systems [9, 10]. This empirical trend can be compared with predictions from the model presented in Equation 5. The model assumption is that the probability of observing a specific mRNA *i* in the sampling process is only determined by its empirical average frequency *f_i_*. It is easy to show [32] that according to this model the probability of not observing a mRNA given the total number of transcripts sampled M is well approximated by

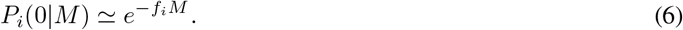

From this expression, we can calculate the expected number of detected different transcripts *h* as

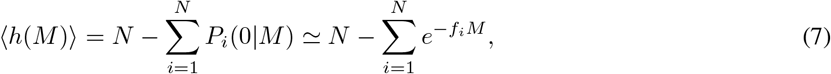

where *N* is the total number of possible mRNAs, given by the number of genes considered in the experiment, which is around 30 * 10^3^. The formula above reproduces well the results of direct simulations of the sampling process (see the Methods section for details) reported as orange dots, and also captures quite accurately the empirical average scaling. Therefore, the observed repertoire of expressed genes in these scRNAseq experiments is on average mostly determined by the sampling process. This trend has to be carefully taken into account in order to reliably estimate the biological variability in transcript repertoires.

A quantitative difference between the empirical average number of expressed genes (blue line in Fig. 3A) and the expectation from sampling (orange line) can be observed. In fact, the sampling model slightly overestimates the empirical trend. In other words, cells typically express a lower number of genes to a higher expression level than expected. This small discrepancy is linked to the statistics of zero values that will be discussed in detail in the following sections.

As illustrated in Fig. 2, two power-law regimes followed by an exponential decay can be sketched for the expression levels. The model can be simplified by exploiting this observation. Instead of considering all the *f_i_* values as free parameters that have to be inferred from data, we can assume the double power-law scaling, with exponents *γ*_1_ and *γ*_2_ estimated by fitting, and the exponential tail for low frequency components. In this case, it can be shown [32] that the expression for *h(M)* simplifies to

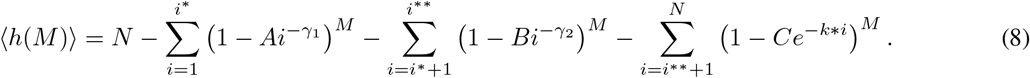

The factors *A, B, C* are defined by imposing normalization and continuity conditions between the three regimes:

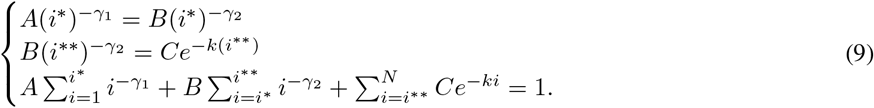

*i*\* is the rank at which the change of power law exponent is estimated, while *i*\*\* is the rank at which the exponential regime starts. This is the theoretical prediction reported as a dashed-red line in Fig. 3A.

If the sampling process is the dominant factor setting the repertoire of observed transcripts, the trend should not depend crucially on the biology of the system in analysis. Indeed, the sublinear scaling is well conserved across different organs as reported in Fig. 3B.

### 3.4 Variability in the repertoire of expressed genes follows Taylor’s law and reveals deviations from a sampling process

As discussed in the previous section, the scaling of the average number of detected genes can be well explained as a result of the sampling process. However, there is substantial variability in the empirical data, i.e., cells with the same total number of UMI can have expression repertoires of largely different sizes. The question is if this variability can be again explained as sampling fluctuations. The model provides a precise prediction for the variance 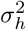 as a function of the average value *〈h〉.* Fig. 4A compares the model prediction of a Poisson scaling (blue dashed line) with the empirical scaling (grey dots) evaluated over all the cells in the MCA dataset in order to have large statistics. The empirical variance displays a power-law scaling with the average vocabulary size that is not compatible with a Poisson scaling. Fitting the empirical scaling with the function *C〈h〉^k^* leads to an exponent *k* = 1.64 ± 0.18. This value is significantly different from the Poisson scaling expected from sampling (*Z* = 3.5) and more compatible with the quadratic scaling (*R*^2^ = 0.94 and *Z* < 2) that has been observed for several other complex systems [11, 33, 34].

**Figure 4:**
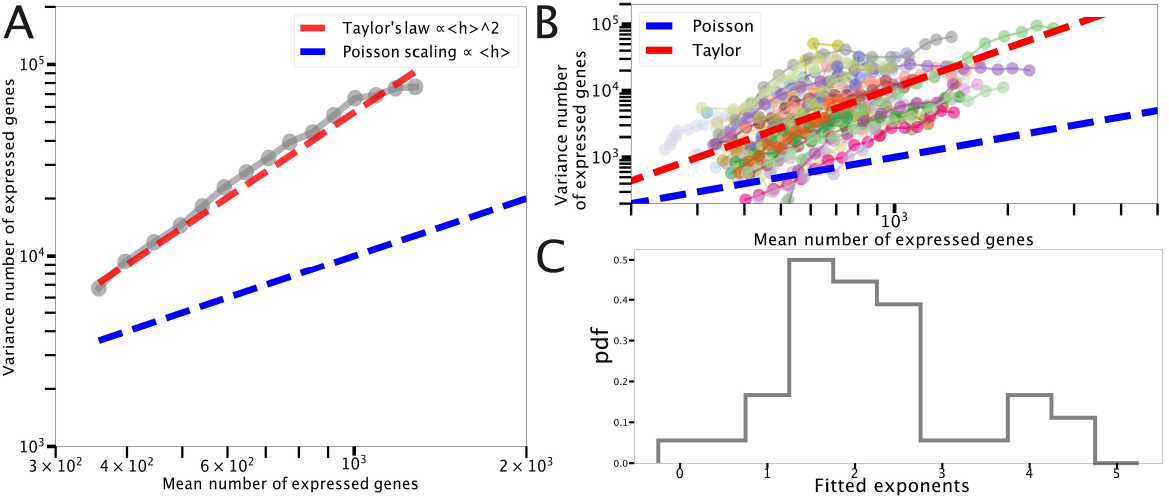
Fluctuation scaling in the number of detected transcripts follows Taylor’s law. **(A)** The variance in the number of measured expressed genes is reported as a function of its average value for all cells in the MCA. Data are compared to a quadratic scaling (red dashed line) and to the Poisson scaling predicted by a sampling process (blue dashed line). **(B)** The fluctuation scaling is conserved by considering separately different organs and tissues. **(C)** Probability density function of the exponents *k* obtained by fitting the curves in panel **(B)** with *C〈h〉^k^*.

Focusing on single cells belonging to the same organ the phenomenology is quite diversified, also due to the reduced cell numbers (Fig. 4B). However, fitting these organ-specific fluctuation curves again with the function *C〈h〉^k^* leads to a distribution of exponents peaked on 2 (Fig. 4C). Although the distribution is quite large, this suggests that an approximately quadratic scaling is an inherent property of the transcriptome diversification and it is not only due to differences between organs.

Interestingly, we have found an emergent statistical law that cannot be explained by the sampling process inherent to RNA sequencing, and that can thus contain information on biological variability. However, this quadratic fluctuation scaling is yet again a common feature of several complex component systems, from linguistics to ecology, known as Taylor’s law [11, 33, 34]. Therefore, a general explanation, which goes beyond the specific properties of expression profiles, could be at the origin of this scaling.

### 3.5 Poisson noise sets the lower-bound and the scaling of gene expression cell-to-cell variability

A commonly analyzed property of cell-to-cell variability in single-cell expression studies is the Coefficient of Variation (*CV* = σ_*n*_/〈*n*〉) of gene expression levels across cells. In particular, the 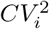 of each gene *i* is often reported as a function of the mean gene expression level in order to identify highly-variable genes at a given average value. In fact, this criterion is often used to reduce the number of features (i.e., genes) to consider in further analysis [35, 36]. We analyzed this fluctuation scaling in the two scRNAseq atlases, and in a bulk RNAseq large-scale experiment (GTEx, see the Methods section) for comparison.

Fig. 5A shows the 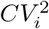 scaling for cells in the MCA. The red dashed line is the analytical prediction (confirmed by simulations) of the expected scaling for a sampling process, which is basically a Poisson scaling. The measured values follow essentially the same scaling, but the observed gene expression fluctuations are larger than the Poisson prediction, which essentially sets the lower bound of measurable variability.

**Figure 5:**
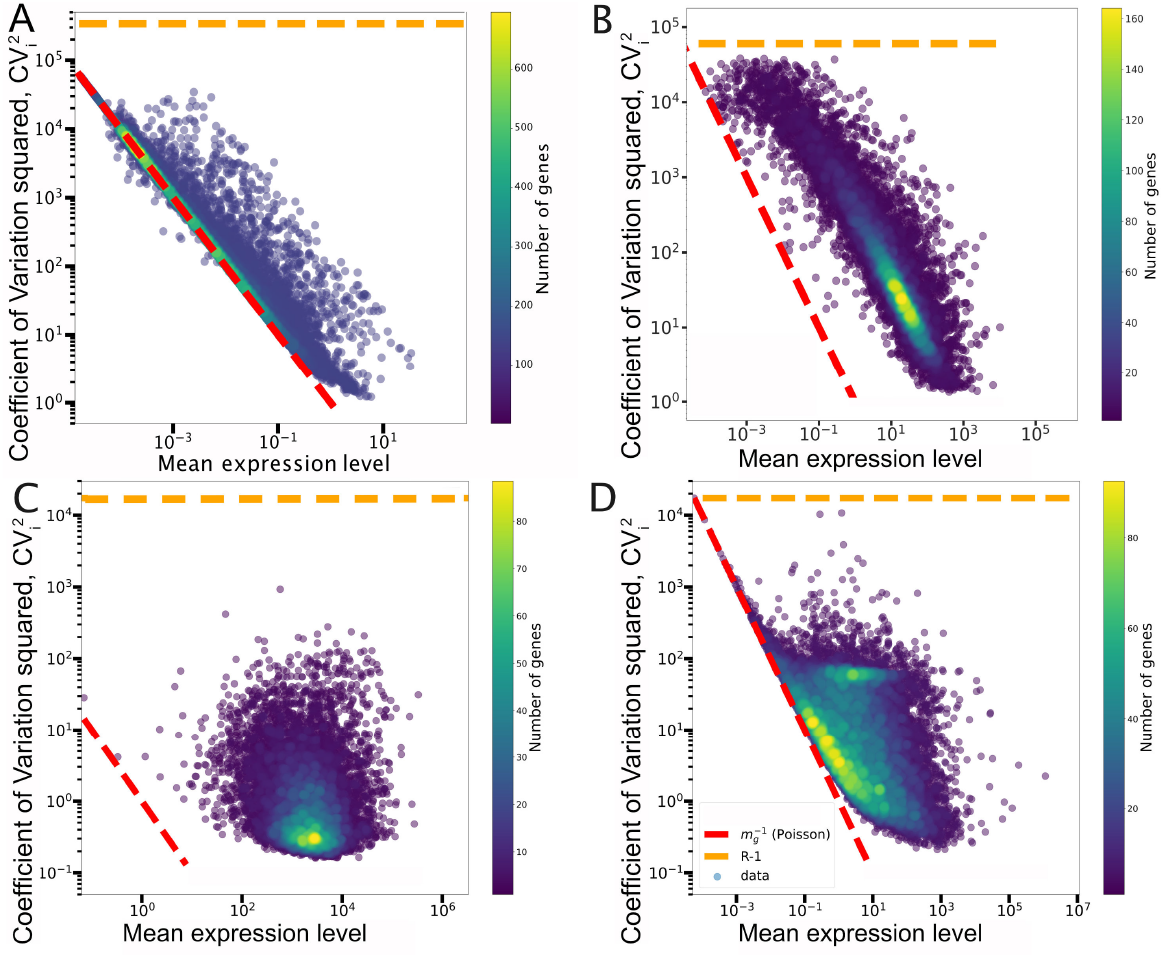
The Coefficient of Variation 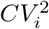 as a function of the average expression. The red dashed lines report the Poisson scaling,i.e., 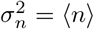. The horizontal orange dashed lines correspond to the maximum possible value of *CV*^2^ = *R* – 1, which is achieved if a gene is expressed in only one cell. The different panels correspond to data from: **(A)** Mouse Cell Atlas; **(B)** Tabula Muris; **(C)** GTEx limited to protein coding genes; **(D)** GTEx limited to non-coding genes. As explained in the Methods section, for the MCA database we considered the UMI counts, while for Tabula Muris and GTEx the raw read counts are reported.

The analogous plot for the Tabula Muris dataset displays the same scaling but with a clear shift. This can be simply explained by the amplification process used before sequencing. In fact, a Poisson random variable multiplied by a constant has a translated *CV^2^*. Consider a random variable *x* with average 〈*x*〉 = *μ* and variance 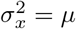. We can now define a new variable *y* = *kx*, where *k* is a constant describing a supposedly constant amplification factor. The mean and variance of *y* are simply given by 〈*y*〉 = *kμ* and 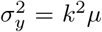. Therefore, the *CV^2^* of the scaled variable *y* is still 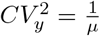 but can be written as a function of 〈*y*〉 as 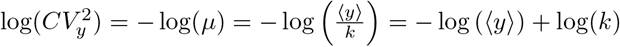.

The *CV^2^* has a natural upper bound that is reached when a gene is expressed in only one cell. If the only not-zero count is *n*, the average expression is *n/R* (where *R* is the number of cells), and the *CV^2^* is *R* – 1, which does not depend on n. This upper bound is reported as a dashed orange line in Fig. 5. Note that the highest variability reported in the Tabula Muris atlas closely approaches the bound.

In bulk RNA sequencing data, the effects of sampling and stochasticity in gene expression should be averaged out by extracting RNA from a large number of cells. In fact, the *CV* profile is radically different if calculated on protein-coding genes in the GTEx dataset (Fig. 5C). The empirical variability is far from the sampling limit and the *CV* seems typically weakly dependent on the average expression level. However, focusing on non-coding genes (Fig. 5D), which are typically lowly expressed, we observe the Poisson trend emerging again. This indicates that, as expected, sampling has to be carefully taken into account when the variability of lowly expressed genes is analyzed even in the context of bulk RNA-seq data.

This analysis shows that the sampling process sets a lower bound to the measured variability in gene expression, and data from single-cell RNA sequencing are generally close to this bound. Clearly, the observed expression variability is not captured by our simple null model which focuses on sampling and thus can only produce Poisson distributions.

Note that a Poisson scaling could also be explained by a simple model of stochastic gene expression in which transcription and degradation are modeled as reactions with constant rates [37, 38]. This description also leads to Poisson distributions for mRNAs and thus to the same scaling of the *CV*. However, transcription can be more complex than a birth-death process, for example it can be characterized by bursts of expression [39]. Models accounting for bursty production naturally lead to overdispersed expression distributions such the Negative Binomial distribution (or its continuous analog Gamma distribution), or even to more complex bimodal distributions [40, 39, 41]. Moreover, the presence of extrinsic noise, i.e., fluctuations in global cellular factors [42], can induce a constant *CV* with respect to the average expression [43], and there could be an additional basal technical noise besides sampling fluctuations in scRNAseq data [19].

Indeed, empirical *CV^2^* values typically present a double scaling, with a Poisson-like dependence for lowly expressed genes (where intrinsic noise and/or sampling effects are relevant) and a constant “floor” noise at higher level of expression both for proteins and for mRNAs [43, 20]. This constant floor noise is also evident in bulk transcriptomic data (Fig. 5), as well as for highly expressed genes in scRNAseq datasets, although the double scaling is more evident for single-cell datasets obtained with the Smart-seq3 technology that will be introduced in a following section. A constant *CV* is precisely equivalent to a fluctuation scaling of the type 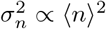. While this functional form is again reminiscent of Taylor’s law [33], gene expression fluctuations are not equivalent to the fluctuations in the number of expressed genes, i.e., the fluctuations around Heaps’ law that we previously described (Fig. 4). In other words, a quadratic scaling of expression fluctuations, for example due to stochastic gene expression, does not necessarily induce a quadratic scaling of the fluctuations around Heaps’ law. A simple model experiment can be used to prove this point. We assume that genes have Gamma-distributed expression levels, with mean values following a Zipf-like law. We also assume that the *CV* of the expression levels is constant, thus a Taylor’s law for expression fluctuations, by appropriately fixing the variance of each Gamma distribution. These Gamma-distributed expression levels are then randomly sampled obtaining a *CV^2^* that displays a double scaling as in many empirical observations (S9A). However, the fluctuations in the number of expressed genes, i.e., the analog of Fig. 4, can still show an approximately Poisson scaling (S9B). Therefore, more realistic models of stochastic gene expression can in principle explain the empirical levels and the scaling properties of expression fluctuations, but they do not necessarily reproduce the Heaps’ law fluctuations.

### 3.6 The statistics of transcript sharing

While the repertoire of expressed genes can be highly cell specific, it is natural to expect a certain degree of overlap between the genes that have to be expressed in different cells. This overlap should depend on the specific gene functions and on the similarity of the cell types in analysis. For example, we intuitively expect a core set of genes, linked to basic cellular functions, to be expressed in essentially every cell. In order to quantify the statistics of the overlaps between the expression profiles of different cells, we analyzed the occurrence distribution. The occurrence *o_i_* of a transcript is defined as the fraction of cells in which it is detected (i.e., it has a non-zero count). Fig. 6A reports the occurrence distribution for cells belonging to a single tissue (the bone marrow in the example). Surprisingly, most of the genes appear to be expressed in very few cells and the number of genes expressed in all cells seems negligible. However, a quantitative comparison with the null model suggests that this is mainly an effect of the sampling process. In fact, given the empirical average expression levels *f_i_*, the sampling model (Eq. 5) gives the expected occurrence for each gene *i* as

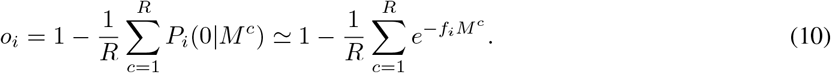

**Figure 6:**
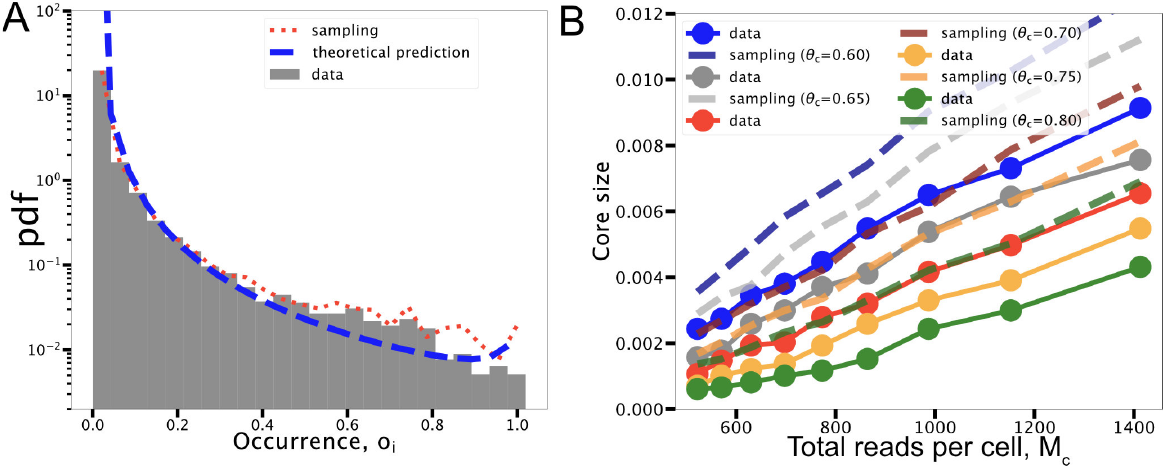
The occurrence distribution of gene expression. **(A)** The probability of observing a mRNA in a fraction of cells is reported for the illustrative example of cells in the bone marrow as profiled in the MCA. **(B)** The empirical fraction of genes expressed in at least *θ* cells (different *θ* values correspond to different curves) is compared with the corresponding predictions of the sampling process.

From this expression, the probability density of the occurrences can be extracted. It is reported as a dotted red line in Fig. 6A and provides a good approximation of the empirical distribution. An equivalent result can be obtained from direct simulations of the sampling process.

As previously shown [6], the occurrence distribution takes a particularly simple functional form if we approximate the distribution of relative expression levels with a single power law (*f_i_* ~ *i*^-*γ*^, with *γ* ≃ −0.8), and we assume that all cells have the same average number of total UMI (M ≃ 1 500 transcripts in this case). The resulting expression is

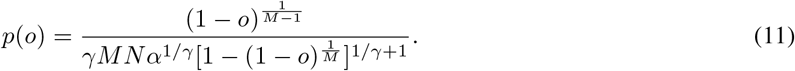

Despite the crude approximations, this analytical prediction (blue dashed line in Fig. 6A) can still reproduce reasonably well the data, and can thus be used in general for an easy first prediction of the effect of sampling noise on mRNA occurrences. While a simple Poisson sampling process can largely explain the shape of the occurrence distribution, there are quantitative differences. In particular, there is a clear difference between the two distributions for high occurrence levels (Fig. 6A): ubiquitously expressed genes, or core genes, seems under-represented in the data. A more detailed comparison can be done by explicitly looking at the core size and how it scales with the total number of sequenced transcripts *M*. The core size *c* can be defined as the fraction of genes expressed in at least a fraction θof the cells in the population, i.e., genes with *o_i_* > *θ*. Considering again the approximation of a power-law distribution of average expression levels, the sampling process predicts a specific scaling for the core size with the sample size *M* [6]. The core size is indeed described by the expression

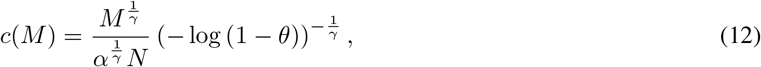

where *α* = ∑_*i*_ *i*^−*γ*^ is a normalization and *γ* ≃ 0.8 is estimated from data. Thus, the scaling is expected to be approximately linear if γ is close to 1. This qualitative prediction is confirmed by empirical data (Fig. 6B). In this plot, core sizes, defined by different values of *θ*, are measured over cells with a different number of detected transcripts *M* (dots in Fig. 6B). The empirical scaling can be compared with direct simulations of the sampling process or with the equivalent numerical integration of Eq. 10 (dashed lines in Fig. 6B). The linear trend described by Eq. 12 is observed in both data and simulations. However, the empirical curves have slightly smaller slopes and they systematically show smaller core sizes. In other words, given the average gene expression levels, there is a smaller than expected number of genes that can be detected in a large fraction of the cell population. The origin of this discrepancy is closely related to the statistics of zero values in scRNAseq datasets that will be addressed in more detail in the next section.

### 3.7 Predicting presence from transcript abundance and the statistics of zero values

The sparsity of scRNAseq data and the possible origins of the detected zero values have been, and still are, an active field of research and debate [14, 13, 44]. Using our simple null model, we can identify what level of data sparsity is expected from sampling only. Moreover, we can isolate genes whose zero statistics is unexpected, and thus possibly linked to biological variability or technical noise not included in our description. As discussed in the previous section, the sampling model provides a prediction for the occurrence *o_i_*, i.e.,the number of cells in which the count is not zero, for each gene (Eq. 10). This prediction can be directly compared with the empirical occurrence as in Fig. 7A and B. The first observation is that the density of points is mostly located on the diagonal, where the number of zero values predicted coincides with the sampling expectation. In fact, the probability density of the differences between the predicted and the empirical occurrences is extremely peaked on zero as reported in Fig. 7C (note that the y-axis is in logarithmic scale). Therefore, the zero values are precisely those expected from sampling for most genes, and this result suggests that complex zero-inflated models, which are often introduced to capture the data sparsity [45,46], are generally not needed. This observation is in line with recent analysis of scRNAseq data based on UMI [14, 20, 46].

**Figure 7:**
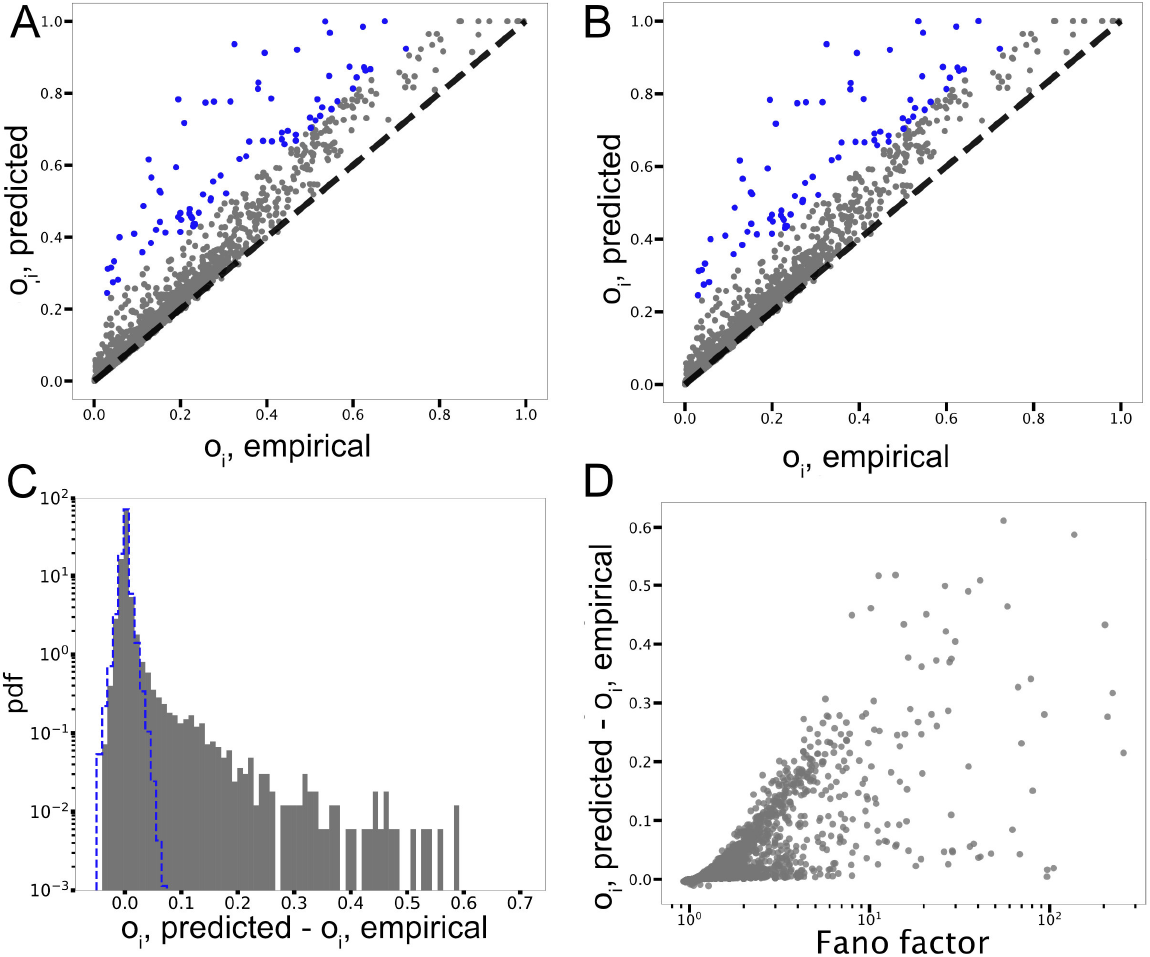
Explaining the empirical occurrences from average expression and sampling. The number of cells in which a transcript is present *o*_*i*, empirical_ is reported with respect to the occurrence predicted by sampling *o*_*i*, predicted_ for **(A)** Bladder and **(B)** Muscle. **(C)** The distribution of *o*_*i*,predicted_ – *o*_*i*,empirical_ is extremely peaked on zero, but with a right tail indicating a higher level of sparsity in empirical data. The blue dashed line represents the distribution of the typical occurrence deviations between sampling realizations, thus providing an estimate of the deviations that are compatible with sampling fluctuations. **(D)**Scatter plot of the relation between the Fano factor 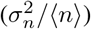 of expression levels and *o*_*i*,predicted_ – *o*_*i*,empirical_ in Muscle as an illustrative example.

However, the fraction of data points that deviates from the diagonal are mostly above it, showing that indeed genes whose zero count statistics is not well described by sampling have an excess of zero values. This leads to a general level of data sparsity that is slightly underestimated by the model. For example, 97%of entries in the Muscle dataset reported in Fig. 7B are zero values, while the sampling process predicts 96%null entries on average. Such a minor discrepancy should not be surprising since our model is an intended over-simplification of the system. For example, we are not considering the inherent variability due to stochasticity in gene expression. We are approximating the true gene expression distributions as delta functions (see Methods section), which become Poisson distributions only through the sampling process. The underestimation of expression variability is explicitly depicted in Fig. 5, where the variance of empirical expression values is often larger than Poisson. The Fano factor or index of dispersion (i.e., 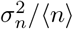) can be used to quantitatively measure how large is the deviation. In our case, it measures how far is a gene expression variability from the sampling prediction. As intuitively expected, the Fano factor is correlated with the difference between the predicted and observed occurrences for each gene (Fig. 7D). However, the scatter plot does not show a clear and simple relation, thus suggesting a complex interplay between expression variability and zero count statistics.

Given its inherent simplifications, the model provides a simple and quantitative way to select the genes whose zero value counts are “atypical” and thus that are potentially interesting for further analysis [47]. This excess of zero values can derive from technical reasons (often called “drop-outs”), from biological variability due to cell-type heterogeneity or eventually from particularly noisy promoters. Subsequent analysis of the selected genes could select the most likely contributions. While we leave this step for future work and specific applications, we propose an illustrative example. We selected the genes with the top values of *o*_*i*,predicted_ – *O*_*i*,empirical_ (depicted in color in Fig. 7A and B) in different organs and performed a GO enrichment analysis. Some categories are over-represented. The presence of enriched GO categories already indicates that if a drop-out phenomenon is present, it is not random across genes. All the significantly enriched categories are reported in S6 of the Supporting Information together with links to the full gene lists. Highly enriched categories could indicate biological signals as well as technical reasons not captured by a simple sampling. For example, the presence of general categories such as “Ribosomal proteins”, “Extracellular exosome” or “Blood microparticle” could be due to the insertion of few zero values at the initial filtering procedure. Ribosomal proteins are expected to be expressed in essentially every cell type at a relatively high level. This is generally the case in the data, since those genes are in the high-rank region of the Zip’s law (Table S2). Therefore, the few empirical zero counts cannot be explained from sampling given the high average expression levels. In this example, the sampling model could be a potentially useful check of the technical procedures.

Note that sampling is a stochastic process, thus the predicted number of zeros for a given gene has a confidence interval determined by sampling fluctuations. These fluctuations can be evaluated using an ensemble of sampling realizations and measuring the distribution of gene occurrences across the ensemble and the typical differences between sampling predictions that can be compared with the empirical ones (blue dashed line in Fig. 7C). For example, the small fraction of genes that are detected in more cells than expected given their average expression (negative values in Fig. 7C) can be explained by sampling fluctuations.

As a further test of the significance of analyzing the deviations from our data-driven null model, we consider few datasets composed by cells from different organs in the MCA, and we focus again on genes whose zero value statistics significantly differ from expectation. The rationale is that genes whose expression is, for example, tissue specific, will typically have expression distributions far from the null model expectation when they are evaluated over cells belonging to different organs. This would also reflect in their atypical zero value statistics. In fact, the genes selected based on their anomalous occurrences (as the coloured dots in Fig. 7) across cells from “Brain” and “Ovary” are significantly enriched in the GO term “myelin sheat” (P-value 6.8 * 10^−16^) which is clearly a tissue specific function that characterize only a fraction of the cells in analysis. The same GO term appears when cells from “Brain” and “Muscle” are considered (P-value of 8.4 * 10^−11^). Considering cells from “Muscle” and “Blood” instead leads to the selection of genes significantly associated to “haptoglobin” and “hemoglobin” (P-values < 10^−6^). The GO term “spermatogenesis” is associated to genes with atypical occurrences in a dataset joining cells from “Ovary” and “Testis”. The full lists of the selected genes in these illustrative examples are reported in the Supplementary Material (S7, S8, S9, S10).

Besides gene selection, a dataset overall deviation from the prediction of the corresponding null model should be related to the inhomogeneity of the cellular transcription programs it includes, thus ultimately to the “complexity” of the cell population analysed. As a preliminary test of this hypothesis, we focus again on the number of detected genes in datasets composed by cells belonging to a different number of organs. The number of organs included can be used as a rough measure of the dataset inhomogeneity. We observe a clear correlation between this number and the statistical deviation from the sampling model (S10). Although we are only focusing on the discrepancy between model and data in the gene occurrences (thus essentially on the Heaps’ law), this result suggests that the null model is a useful tool to measure and quantify intrinsic properties of the dataset.

### 3.8 Checking the robustness of the statistical laws

As a further test of the robust emergence of the described scaling laws, we analyzed two additional datasets of cells profiled with the recently introduced protocol Smart-seq3 [18]. The Smart-seq3 protocol combines high sensitivity with the use of UMI, providing reliable molecule counts and data matrices typically with a lower degree of sparsity. Despite the differences in the protocol and in the cell types considered, the same phenomenological laws reported for the large-scale mouse cell atlas are clearly observable. Also in this case, the general trends can be framed in our analytical framework and partially explained by a sampling process. Fig. 8 shows some of these laws for the example of a HEK cell line, while the analogous results for mouse fibroblasts are reported in S11. The rank-plot of the average expression levels is again a Zipf-like law characterized by three clearly distinguishable regimes with a central power law scaling with exponent close to −1 (Fig. 8A). Interestingly, the UMI-based datasets present a fitted exponent with values very close to the classic −1 (Table S2). The trend predicted by the sampling process for Heaps’ law (Fig. 8B) is compatible with the empirical number of detected transcripts, with a slight sampling overestimation that is linked to the zero-value statistics. The cell-to-cell variability in expression levels shows a Poisson scaling for lowly expressed genes. However, given the higher sensitivity of the Smart-seq3 protocol, the regime of approximately constant *CV*, which was observed for bulk data (Fig. 5), is now detectable for highly expressed genes, where sampling effects are less dominant. Interestingly, this double scaling of the *CV*^2^ is analogous to the one reported for protein fluctuations in large-scale single-cell experiments based on fluorescence [43, 48].

**Figure 8:**
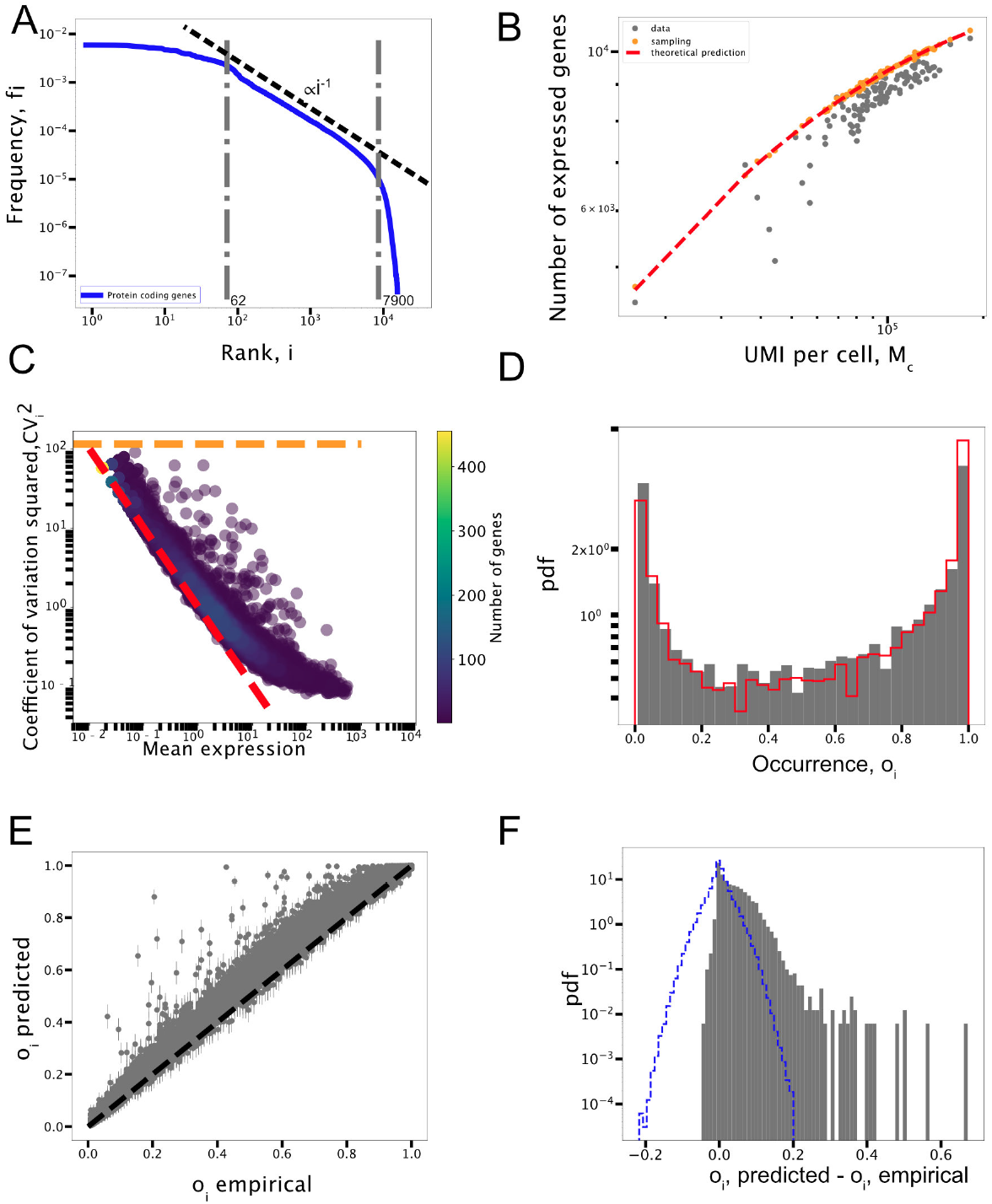
Emergent statistical laws from a HEK cell line profiled with Smart-seq3. (**A**) Zipf’s law with the three scaling regimes. (**B**) Heaps’ law: number of detected transcripts as a function of the total number of UMI per cell. (**C**) Expression variability *CV*^2^as a function of the average expression level for all detected transcripts. (**D**) The empirical transcript occurrence distribution compared with the model expectation (red continuous line). (**E**) The relation between the empirical occurrences *o*_*i*,empirical_ and the expected values from the sampling model *o*_*i*,predicted_. Error bars represent the variability (one sigma) between different realizations of the sampling process. (**F**) Distribution of *o*_*i*,predicted_ – *o*_*i*,empirical_. As in Fig. 7C, the blue dashed line identifies the differences compatible with sampling fluctuations.

In the cell line considered for Fig. 8, the empirical fraction of zero values is 45%, while the sampling expectation is 41%. Therefore, the zero statistics is again largely explained by sampling effects. Indeed, the occurrence distribution is largely recapitulated by the sampling model (Fig. 8D). Thanks to the protocol higher sensitivity, leading to lower data sparsity and larger realization sizes *M*, occurrences clearly display the typical U-shaped distribution that robustly emerges in several complex component systems [6]. The observable deviations from the model only derive from the small fraction of transcripts that present more zero values than expected (Fig. 8E and F), in perfect analogy with our results for the MCA. Typical occurrence fluctuations only due to sampling (blue line in Fig. 8D) are expected to be larger for this dataset with respect to the MCA because of the lower number of cells profiled (i.e., 117 cells for the HEK cell line). For this reason, we considered the fibroblasts dataset which contains 369 cells to select genes with more zeros than expected and perform GO enrichment analysis. The results show cell-cycle and cell-division related terms as enriched (S11). Indeed, in a non-synchronized proliferative cell line as the one in analysis, we should expect non trivial expression distributions for genes related to the cell-cycle progression [18].

In conclusion, the presented statistical laws seem a robust emergent property of single-cell RNAseq data. The proposed mathematical framework provides an explanation for most of the general trends, and thus can be a useful simple null model to identify significant deviations of biological or technical origin.

## 4 Discussion

The identification of statistical laws is a key step in designing effective descriptions of complex systems [49]. Leveraging on large-scale regularities, phenomenological models can be built, in the spirit of statistical physics, to capture relevant system properties without focusing on a detailed description of the high number of degrees of freedom. For example, the presence of quantitative empirical laws in cell composition of fast-growing bacteria has lead to simple models of cell physiology that can explain several large-scale gene expression patterns using just few key parameters such as the growth rate [50]. Analogously, the emergence of different cell identities and their organization in tissues and organs is driven at the molecular level by the complex orchestration of the expression of large sets of genes. However, simple coarse-grained descriptions can be hopefully extracted without resorting to all the molecular details. As a first step in this direction, we identified several statistical laws emerging in single-cell transcriptomic profiles using large-scale expression atlases in mouse. Strikingly, analogous laws are ubiquitously found in different complex component systems from linguistics to ecology [9, 11, 51, 6].

An additional complication of scRNAseq data is the presence of a sampling process inherent to the experimental technique. Therefore, the observed expression statistics is due to a combination of natural cell-to-cell variability and stochastic sampling. We focused on modelling the sampling process given a basic system property, which is the specific average heterogeneity of gene expression levels described by the classic Zipf’s law. This law is apparently a hallmark of several component systems [22, 52]. It has been previously reported for gene expression at the population level [24], but we showed that it is an intrinsic property of single cells robustly emerging in different datasets.

The proposed simple model essentially neglects biological expression fluctuations and tests what can be explained from sampling only. In this framework, there is a natural predicted connection between Zipf’s law and other statistical regularities such as Heaps’ law and the U-shaped statistics of shared components [6, 32]. We first showed that indeed these additional empirical laws emerge in transcriptomic data, and second that they can be well explained as consequences of stochastic sampling. This result suggests that downstream analysis typically performed on these datasets, such as clustering to identify cell types or expression fold-change analysis, have to carefully take into account sampling and the statistical regularities it generates.

However, we identified some clear deviations from sampling predictions. Specifically, the empirical variability in the cell expression repertoires, captured by the fluctuation scaling of the Heaps’ law, cannot be reproduced by the model. This result could conceal a biological motivation linked to the differentiation of expression programs in different cell types. However, the very same scaling is a recurrent feature of several complex systems, often called Taylor’s law [33, 34, 11], suggesting a more general mechanism behind its emergence.

This fluctuation scaling is closely linked to the statistics of zero values, which is a central theme in scRNAseq data [14, 13, 46, 44]. In this regard, we first showed that the vast majority of zero counts in the data can be simply explained as a sampling effect. Therefore, there is not a clear indication that complex (and parameter rich) models, such as zero-inflated models, are needed to capture the technical noise [46]. Despite this general trend, the model is a tool to identify specific deviations. A possible application of a data-driven null model that captures general statistical properties of scRNAseq data is to focus on the empirical deviations from its expectation. For example, in several datasets a fraction of genes are expressed in less cells than expected from sampling (i.e., they have an excess of zero counts). We discussed few examples in which the atypical expression distribution of these genes can be explained by their associated biological functions, suggesting the potential use of our null model for gene selection. Interestingly, a similar approach to identify informative genes by comparing their expression variability to a simple random null model was recently proposed [53], precisely leveraging on the analogy between linguistics and transcriptomic data.

The sampling model provides essentially a lower bound for the number of zero values of a transcript given its average expression level. This should be expected since the expression variability in the model only derives from sampling, and thus does not match the typical *CV* values observed. The subsequent step, which we leave for future work, would be to include progressively more realistic models of the stochastic process of gene expression in order to leave out from the description only the cell-to-cell variability coming from the diversity of gene expression programs in the cell population. The price of increasing the complexity of null models is that more parameters have to be introduced and inferred from data.

Statistical laws have also been observed and studied using complex systems approaches at the basic level of DNA sequences [54, 55, 56]. The possibility of a link between general statistical properties at the nucleotide level and the emerging laws for expression patterns here described is captivating, but still to be explored.

Finally, this work adds single-cell transcriptomics to the list of complex component systems displaying statistical laws that are seemingly universal. However, the specificity of transcriptomic data can provide useful indications and constraints to the research of general models and principles behind these laws. Many of the models proposed for the emergence of Zipf’s law in component systems are based on a stochastic growth process. Some examples are classic models based on the Yule-Simon process, on the Chinese Restaurant Process, on Polya urns or on the preferential attachment principle [10, 57, 26]. Basically, these generative mechanisms assume a reuse or duplication of existing components proportional to their current frequencies, and a parallel innovation process that adds new components from a vocabulary. These simple ingredients (with some general prescriptions) are sufficient to reproduce Zipf’s law and the average sublinear scaling of Heaps’ law. The recently proposed sample-space-reducing process can be also be ascribed to this class of stochastic growth models [58, 32]. The description can be appropriate for texts that are generated by the writing process through the progressive addition of words, or for the evolutionary processes that shaped genome composition by duplicating, removing or discovering/transferring new genes. However, the composition of a cell transcriptome is not naturally described by this type of processes, since single transcripts are not progressively added in the cell.

Few alternative compelling mechanisms have been proposed that do not rely on a growth process and could thus apply to the case of transcriptomic data. A possibility is that components have specific networks of dependencies and that these functional relations determine their co-occurence in a realization [59, 8, 21]. In the transcriptomics case, this would translate in an underlying unobserved network of gene-gene dependencies for example due to correlated functions. Models based on this network assumption can generate Zipf’s and Heaps’ laws [21]. Even more generally, power-law distributions can naturally arise if the observed variables (i.e., the expression levels) are affected by fluctuating latent variables that governs the hidden structure behind the data [60, 61]. Gene expression is controlled by several latent factors that defines the state of the cell and are not directly observed in transcriptomic datasets. These latent factors can be highly variable and thus can naturally generate Zipf’s law under certain quite general conditions. One simple example of a hidden variable is the physiological state of the cell, for example described by the growth rate, which is known to strongly influence the gene expression program and the behaviour of different genetic circuits [62, 63]. Analogously, the cell-cycle stage, the cell type or the slowly varying concentration of key enzymes can in principle represent latent variables that have a specific variability in our system and affect gene expression.

## Author Contributions

MO designed of the research. SL and FV performed the analyses and built the figures. MO, SL, FV wrote the article. MO, SL, FV, AM, AS and MC read and edited the manuscript. All authors contributed to the article and approved the submitted version.

## Acknowledgments

We would like to thank Jacopo Grilli, Matteo Cereda and Sarah Perrone for useful discussions.

## Data Availability Statement

The notebooks and the code to reproduce this study can be found in the GitHub repository at https://github.com/BioPhys-Turin/Emergent_Laws_in_scRNA-seq_Data.

## Supporting Information

**Figure S1:**
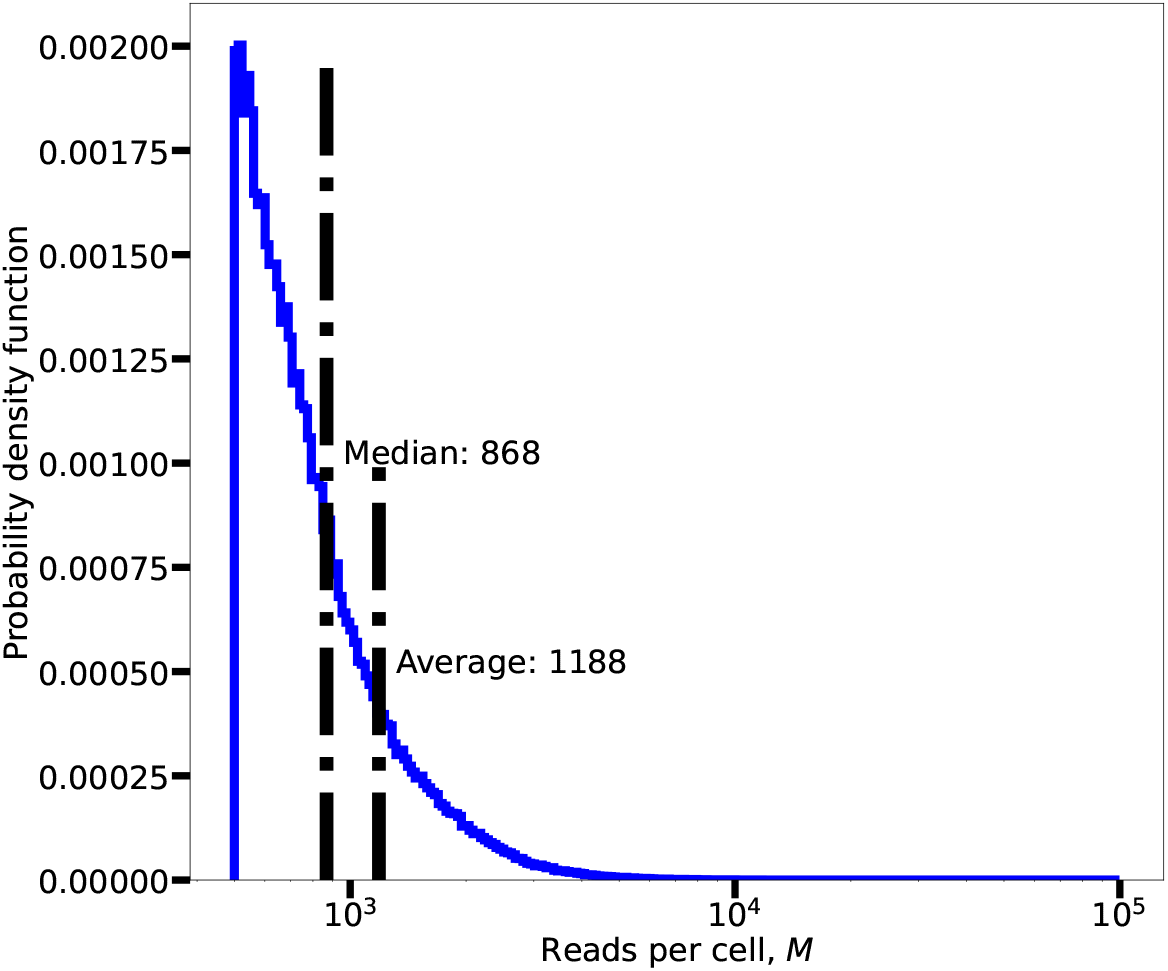
Histogram of the transcriptome sizes in the Mouse Cell Atlas. The transcriptome size *M_j_* represent the total number of UMI detected from cell *j*. Mean and median values are reported for reference.

**Figure S2:**
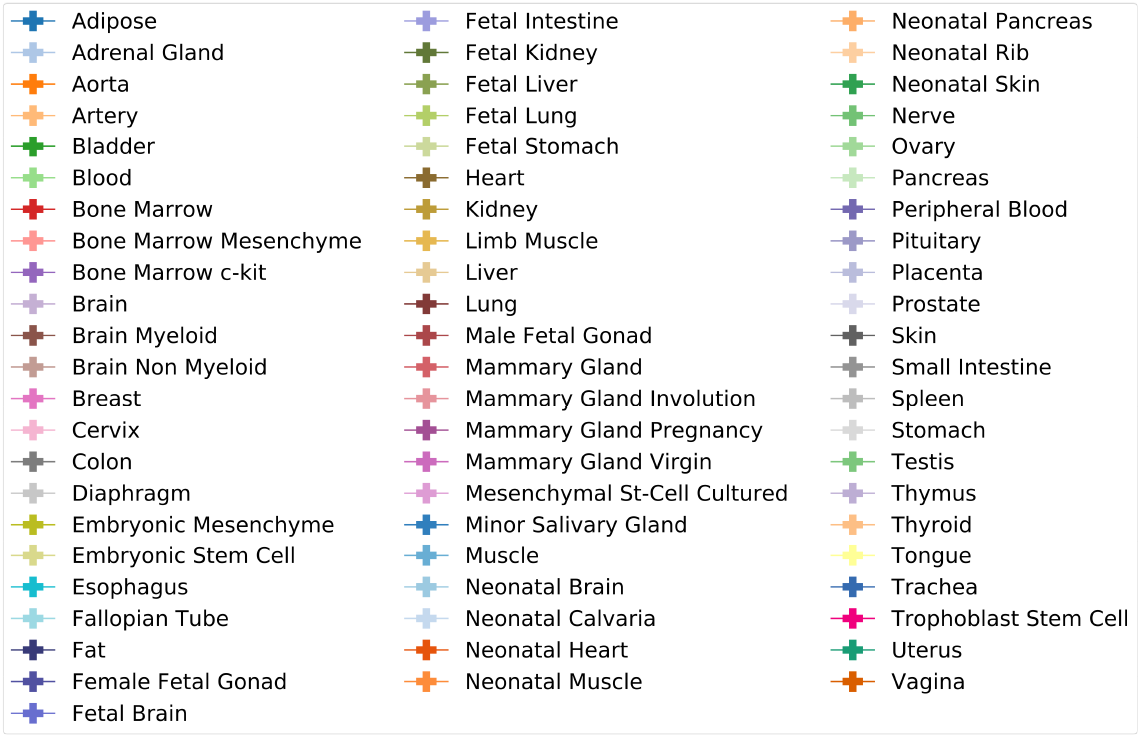
Color legend for the different organs and tissues. All the plots with multiple organs or tissues in the main text refers to this legend.

**Figure S3:**
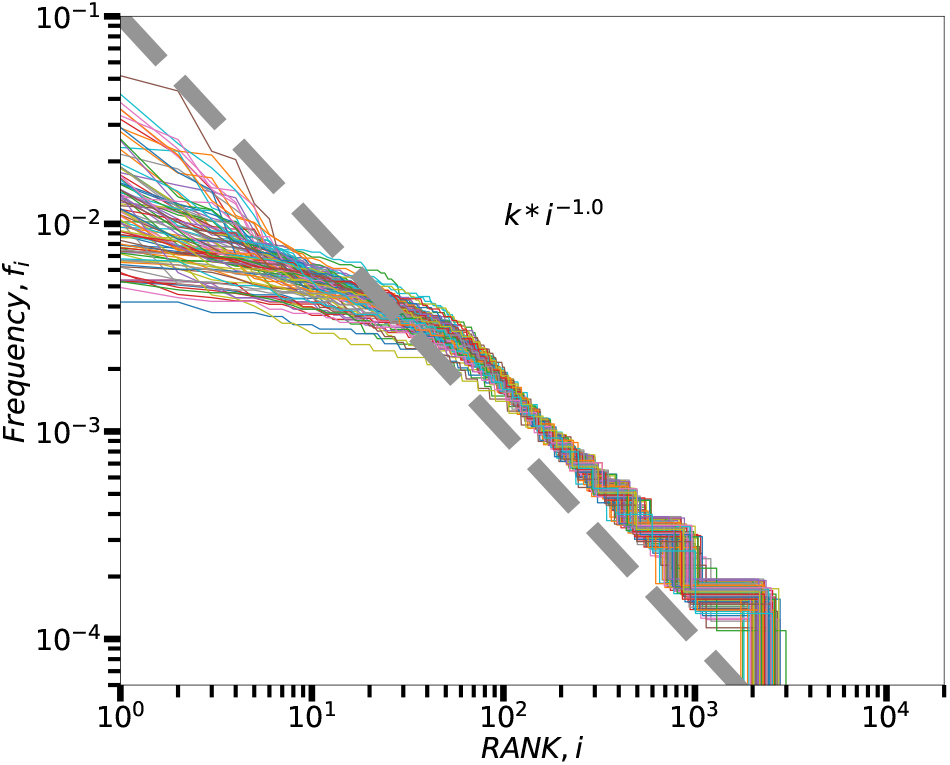
Single-cell Zipf’s law from the Mouse Cell Atlas. Rank plot of the expression levels of the 100 most sequenced cells from the bone marrow in the Mouse Cell Atlas.

**Figure S4:**
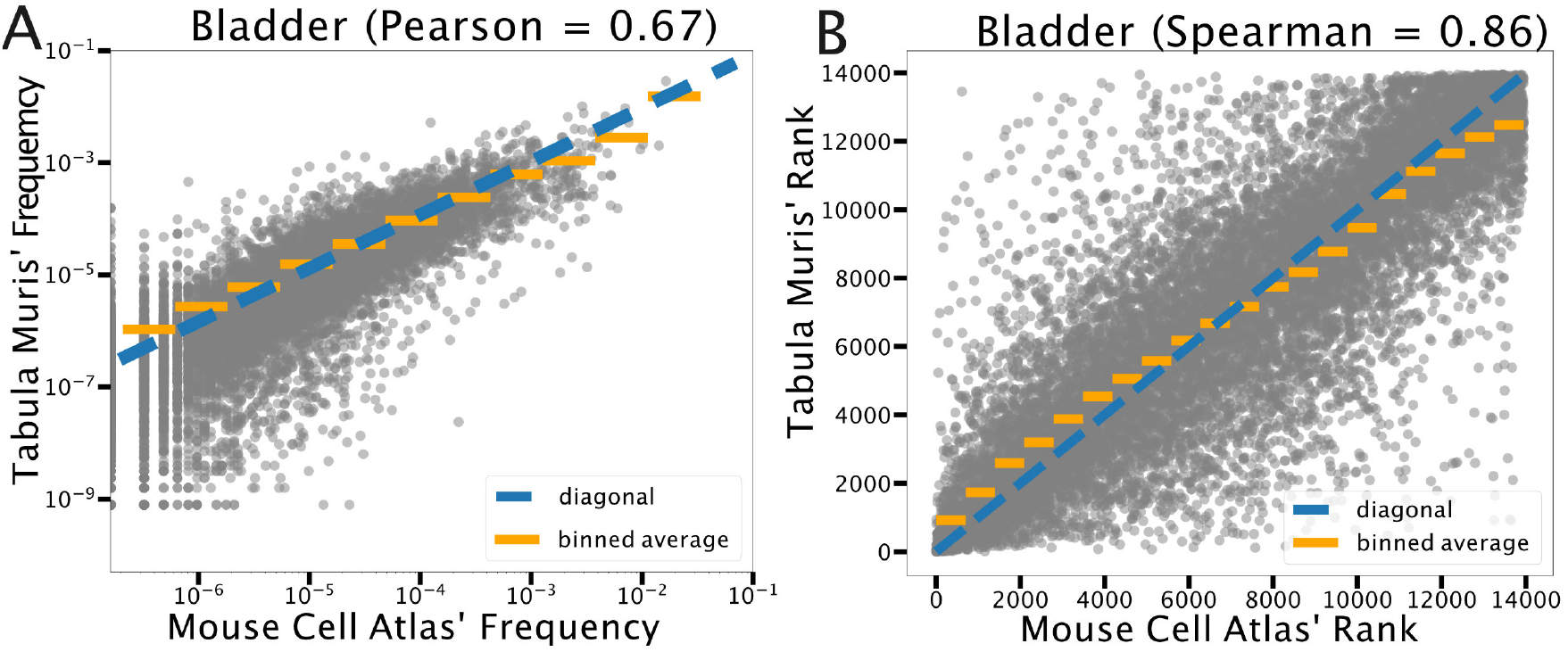
Conservation of expression levels and corresponding ranking across the two different cell atlases. **(A)** Scatter plot of the relative abundances of gene transcripts as reported in the Mouse Cell Atlas with respect to the Tabula Muris compendium for one illustrative organ (equivalent results are obtained looking at the other organs). **(B)** The correlation between the ranking of the relative abundances in the two atlases. Both measures indicate a clear correlation but the variability is substantial.

**Figure S5:**
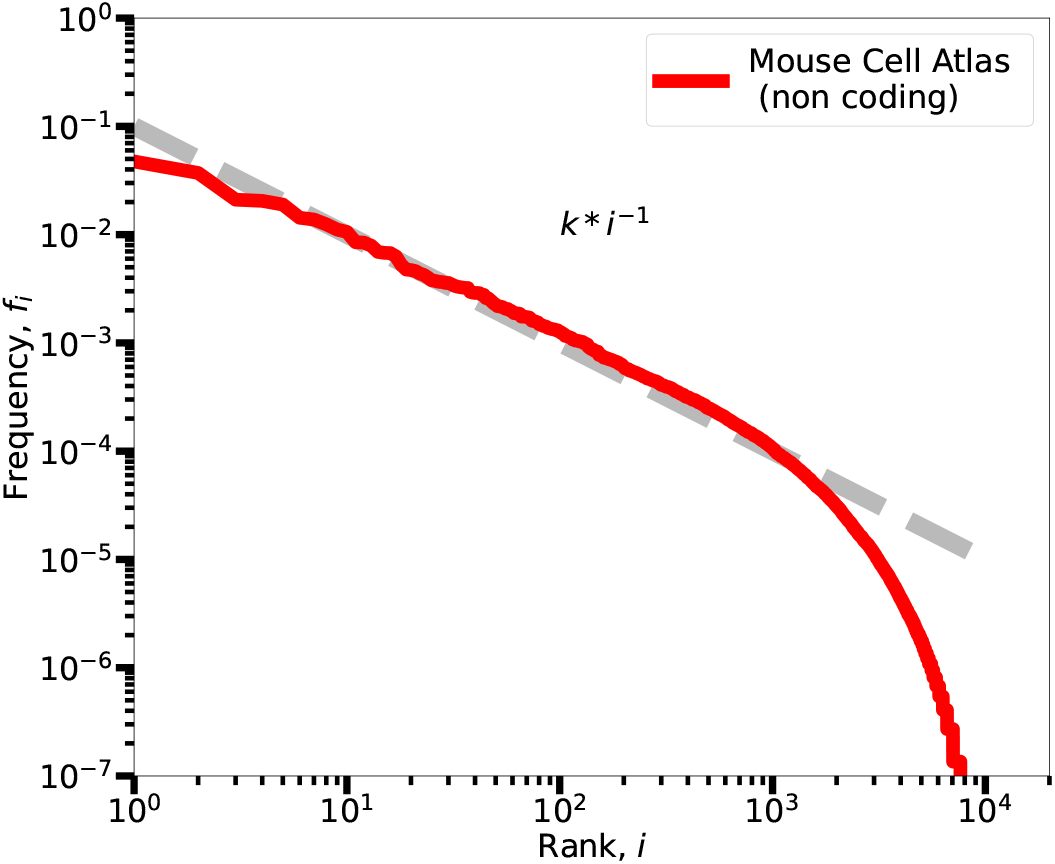
Zipf’s law for non-coding genes. Expression frequencies *f_i_* of non-coding genes from the Mouse Cell Atlas are reported in a rank plot. The power-law scaling followed by an exponential tail is compatible with the Zipf’s law identified for coding genes.

**Figure S6:**
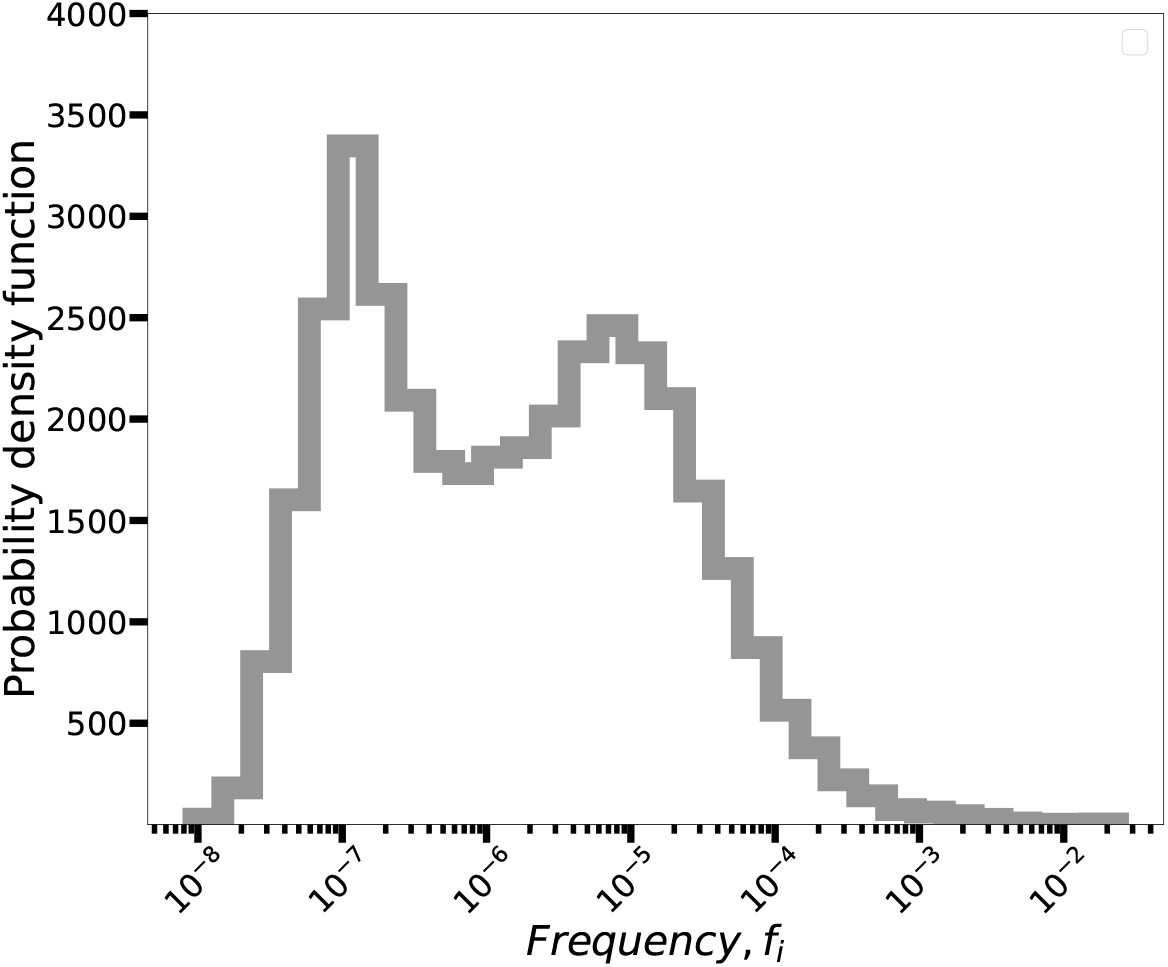
Bimodality in the probability density function of expression levels. We report the histogram of the estimated average transcript frequencies as 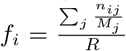. The distribution has a bimodal shape with two maximum values at around 10 ^−7^ and 10 ^−5^. These values correspond to the second and the third regimes of the Zipf’s law we discussed.

**Figure S7:**
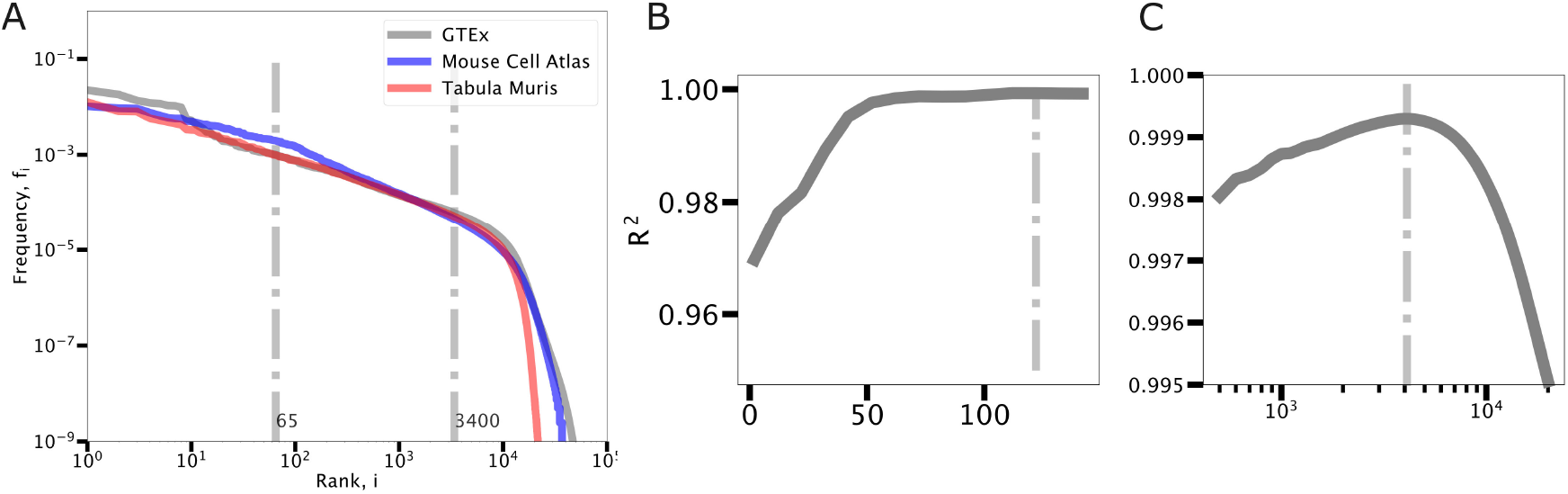
Procedure for the identification of the three regimes in Zipf’s law. **(A)** The average rank plot for each dataset and the average identified bounds. We performed a power-law fit of the frequencies inside a window of ranks with boundaries with variable position. In particular, the left bound was changed from 2 to 122. For each left bound position, the right bound moved between 500 and 20000. For each window position, we fitted with a power-law function and estimated the coefficient of determination 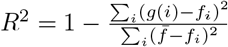. *g(i)* = *A* * *i*^−*γ*2^ is the power-law function of rank *i* obtained by fitting, while *f_i_* are the empirical frequencies with average 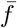. This coefficient is reported in panel **(B)** for the MCA example as we move the left boundary while the right boundary is fixed at its maximum possible value, while in panel **(C)** is reported for different position of the right boundary. The boundaries providing the larger *R*^2^(dot-dashed lines) were selected to identify the three regimes. With fixed boundaries, we finally fitted with another power law the first regime (high frequencies) and with an exponential function *g(i)* = *C* * *e* ^−*γ*_3_\**i*^ the low frequency tail. The left bound was changed from 2 to 122 with a 10 step. Given a left bound, the right bound changed between 500 and 20000 with step of 100. For each dataset we selected the bounds that led to the greater *R*^2^.

**Figure S8:**
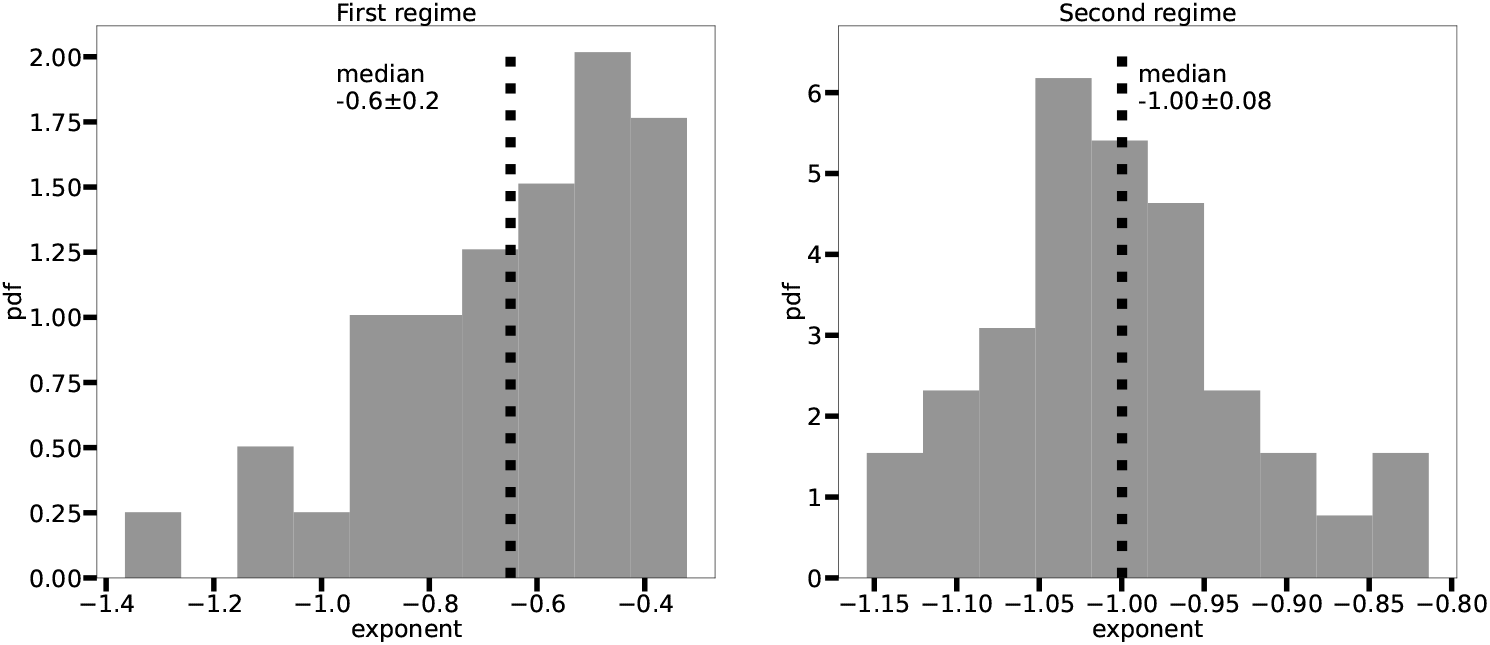
Parameters of the three-regime Zipf’s law for single organs in the MCA. The distributions of the exponents of the power laws fitted in the first two regimes are reported with the corresponding median values. Despite the variability, the median values confirm the presence of a central region with a a trend compatible with the classic Zipf’s exponent -1 after a “flatter” first regime. The median values of the exponents estimated from single tissues are compatible with the ones obtained by fitting the whole dataset.

**Table S1:** Boundaries separating the three regimes of the Zipf’s law. The rank boundaries separating the three regimes obtained by optimizing *R*^2^ are reported for each dataset analysed.

| Dataset | Left Bound | Right bound |
| --- | --- | --- |
| Mouse Cell Atlas | 122 | 4100 |
| Tabula Muris | 12 | 1000 |
| GTEX | 62 | 5100 |
| Average | 65 | 3400 |
| HEK (SmartSeq3 profiled) | 62 | 7900 |
| Fibroblasts (SmartSeq3 profiled) | 82 | 1100 |

**Table S2:** Parameters of the three-regime Zipf’s law in different datasets. Parameters of the first, second and third regimes fit functions (two power-laws and an exponential) *A* * *i*^−*γ*_1_^, *B* * *i*^−*γ*_2_^, *C* * *e*^−*k*\**i*^ in different datasets.

|  | MCA | TM | GTEX | HEK | Fibroblasts |
| --- | --- | --- | --- | --- | --- |
| first regime | $0.515 \pm 0.004$ | $3.4 \pm 0.2$ | $0.73 \pm 0.03$ | $0.35 \pm 0.006$ | $0.32 \pm 0.01$ |
| second regime | $0.9856 \pm 0.0005$ | $0.7588 \pm 0.0006$ | $0.7179 \pm 0.0004$ | $0.98 \pm 0.0009$ | $0.8699 \pm 0.0009$ |
| third regime | $0.00022931 \pm 8 \cdot 10^{-8}$ | $0.0002797 \pm 3 \cdot 10^{-7}$ | $0.0002298 \pm 1 \cdot 10^{-7}$ | $0.00027 \pm 1 \cdot 10^{-7}$ | $0.0003492 \pm 3 \cdot 10^{-7}$ |

**Figure S9:**
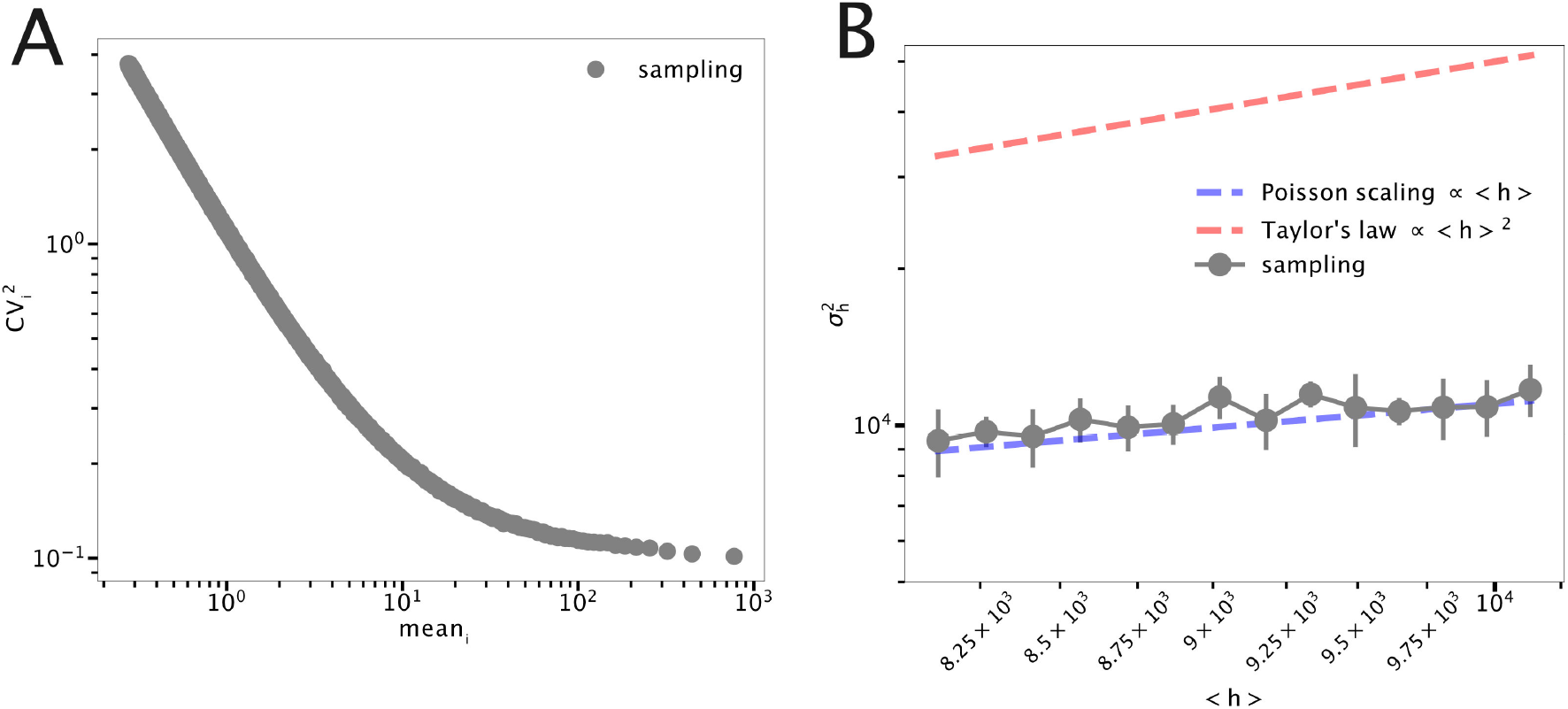
Non-trivial relation between expression fluctuations and variability in the number of detected transcripts. This Figure investigates the link between the fluctuation scaling in expression and the fluctuation scaling around Heaps’s law in an illustrative model. We assume that all gene transcripts are distributed as Gamma distributions with a constant *CV* (thus with 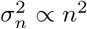). We used the value *CV*^2^ = 0.1 analogous to the empirical value measured for HEK cells (Fig. 8). We considered 20 * 10^3^different transcripts with mean expression values distributed with a Zipf’s law. 2000 artificial cells were then obtained by sampling a total number of transcripts ranging from 20 * 10^3^to 30 * 10^3^. **(A)**The 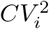 shows a double scaling: for lowly expressed genes we observe the Poisson scaling induced by sampling, while gene expressed highly enough display the floor noise due to the quadratic scaling 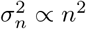. **(B)**A dataset built using this model assumptions presents a Heaps’ law analogous to the empirical ones reported in Fig. 4. We can analyze the fluctuations (i.e., 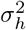) in the number of expressed genes as a function of the average value 〈h〉. These fluctuations do not show the quadratic scaling (in red) that we imposed to fluctuations in expression, and that can instead be approximately observed in empirical datasets (Fig. 4). Therefore, more complex models of gene expression leading to overdispersed mRNA distributions even with a quadratic fluctuation scaling (i.e., a constant *CV*) are not necessarily sufficient to explain the Taylor’s law observed across datasets for the number of expressed genes.

**Figure S10:**
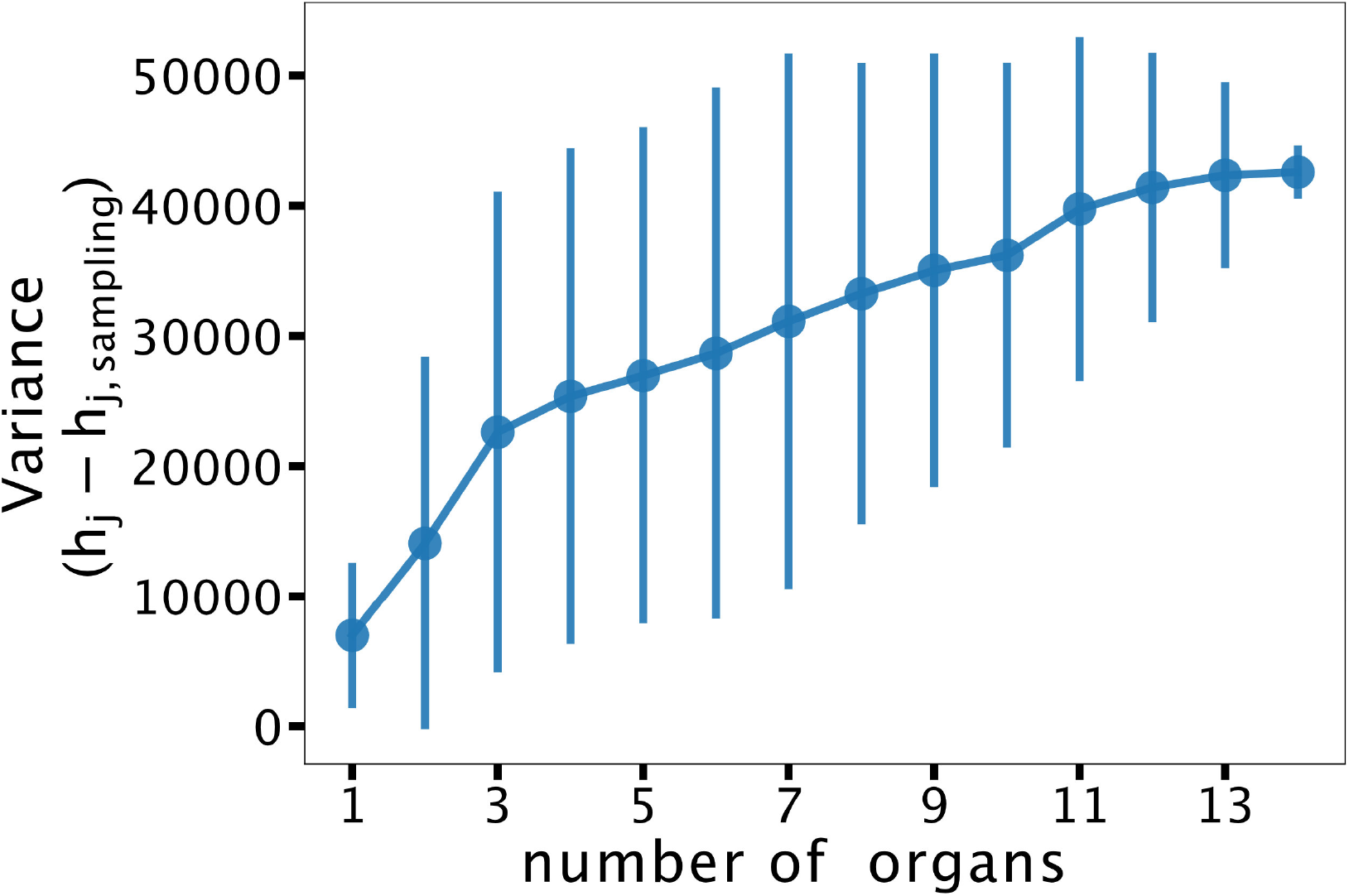
Correlation between dataset complexity and its global deviation from the sampling model in the number of detected transcripts. We built datasets composed by cells belonging to an increasing number of organs by adding 200 randomly chosen cells from a different organ in the MCA. For each dataset we compared the number of expressed genes *h_j_* in each cell *j* with the corresponding typical value *h_j,sampling_* obtained by sampling the same number of total transcripts. We then used the variance of *h_j_* – *h_j,sampling_* across cells to have a global measure of the deviation from the sampling model. A dataset with cells from a certain number of organs can be constructed by assembling different organs. Therefore, we considered 50 different random permutations of the organs in MCA, and this provides the error bars corresponding to a standard deviation in the plot. Despite the large variability between different organ assemblies, there is a clear correlation (Pearson correlation is 0.96). Although in this example we focused on the discrepancy between model and data in the Heaps’ law, this result suggests that the null model is a useful tool to measure and quantify intrinsic properties of the dataset.

**Figure S11:**
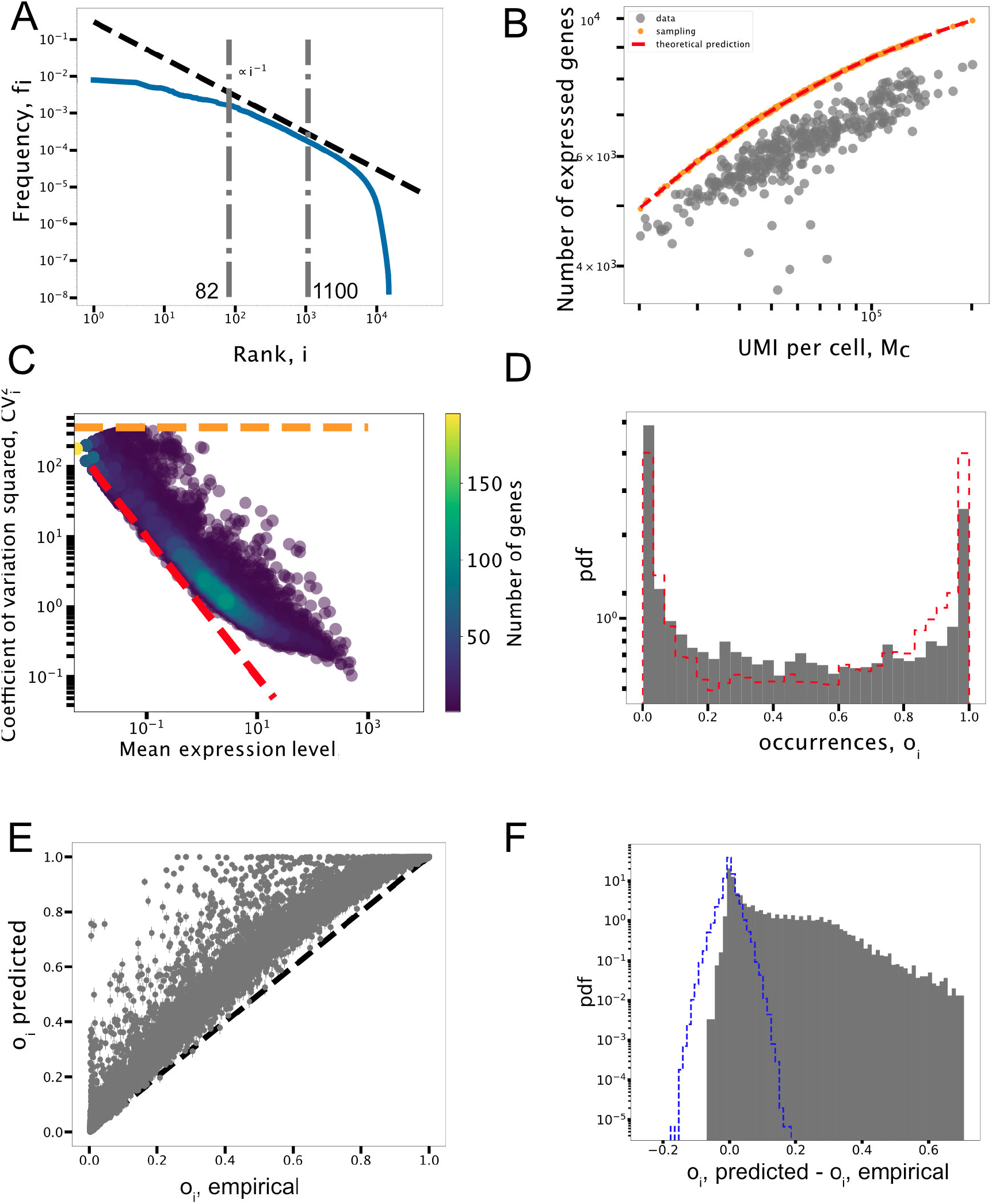
Statistical laws from fibroblasts cells profiled with Smart-seq3 protocol. The Figure is the equivalent of Figure 8 in the main text. (**A**) Zipf’s law. (**B**) Heaps’ law. (**C**) *CV*^2^ scaling with the average expression level. (**D**) Transcript occurrence distribution compared with the model expectation. (**E**)The comparison of predicted occurrences *O*_*i*,predicted_ versus empirical ones *o*_*i*,empirical_. (**F**) Distribution of *o*_*i*,Predicted_ – *o*_*i*,empirical_ and the corresponding distribution from sampling variability.

## Enrichment analyses

We reported in this section all the terms and the Benjamini corrected P-values of gene enrichment analyses as discussed in the main text. They were performed using DAVID and cross-checked using Metascape.

---

**Table S3:** Most enriched GO categories for highly expressed genes in the first regime. Genes with high levels of expression belonging to the first regime of the Zipf’s law in different organs were selected for the analysis. The GO reported here were obtained using DAVID and they are quite similar to those ones found using an alternative tool, Metascape, which are reported in Tab S4. The complete lists of the first regime genes are available at https://github.com/BioPhys-Turin/Emergent_Laws_in_scRNA-seq_Data/tree/main/MouseCellAtlas/firstRegime_lists.

| Organ | GO term description | Benjamini |
| --- | --- | --- |
| Bladder | ribosome | $4.4 * 10^{-100}$ |
| | structural constituent of ribosome | $1.1 * 10^{-97}$ |
| | translation | $7.9 * 10^{-78}$ |
| | intracellular ribonucleoprotein complex | $3.9 * 10^{-68}$ |
| Brain | myelin sheath | $1.9 * 10^{-29}$ |
| | extracellular exosome | $4.6 * 10^{-15}$ |
| | extracellular space | $4.3 * 10^{-7}$ |
| | substantia nigra development | $6 * 10^{-5}$ |
| Kidney | extracellular exome | $3.2 * 10^{-6}$ |
| | mitochondrial inner membrane | $1.3 * 10^{-22}$ |
| | structural constituent of ribosome | $3.9 * 10^{-22}$ |
| Liver | ribosome | $1.7 * 10^{-39}$ |
| | cytosolic small ribosomal subunit | $1.4 * 10^{-38}$ |
| | structural constituent of ribosome | $5.8 * 10^{-38}$ |
| | intracellular ribonucleoprotein complex | $6.9 * 10^{-29}$ |
| Muscle | structural constituent of ribosome | $1.8 * 10^{-61}$ |
| | ribosome | $1.9 * 10^{-60}$ |
| | translation | $3.8 * 10^{-45}$ |
| | intracellular ribonucleoprotein complex | $1.9 * 10^{-43}$ |
| | cytosolic small ribosomal subunit | $1.43 * 10^{-38}$ |
| Pancreas | ribosome | $3 * 10^{-56}$ |
| | structural constituent of ribosome | $2.6 * 10^{-53}$ |
| | translation | $1.7 * 10^{-42}$ |
| | intracellular ribonucleoprotein complex | $1.8 * 10^{-39}$ |

**Table S4:**
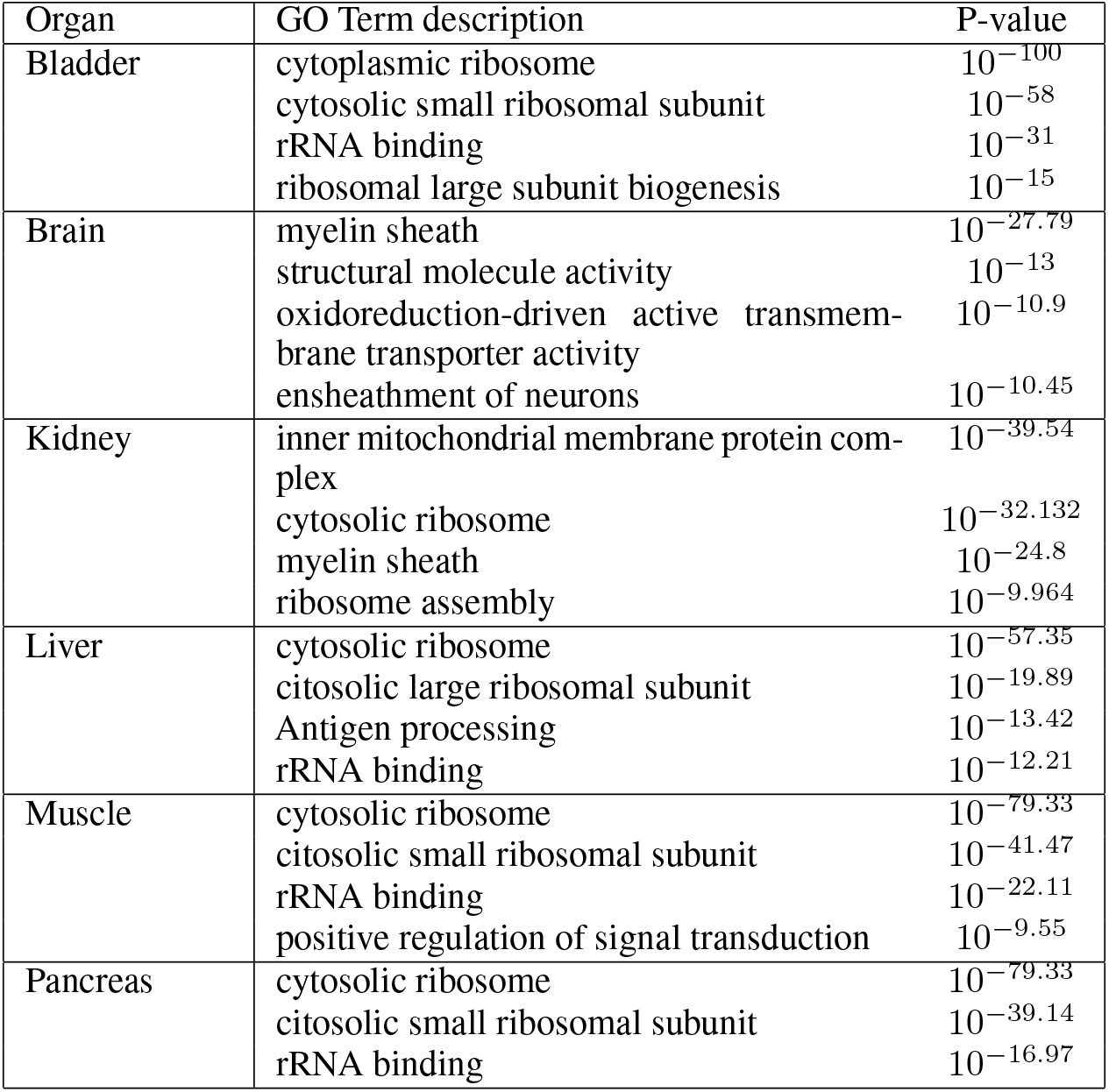
Most enriched GO categories for high expressed genes, using Metascape. We analysed the GO reported in Table S3 using an alternative tool, Metascape. Genes with high levels of expression belonging to the first regime of the Zipf’s law in different organs were selected for the analysis.

**Table S5:** Common genes in the first regime. Here we report the list of the first regime genes common to the 70%of the organs. GO analysis, performed both using DAVID and Metascape, returned “Structural constituent of ribosome” and “Ribosome” as the most enriched categories, with P-values < 10^−43^.

|  |  |  |  |
| --- | --- | --- | --- |
| Calm1 | Rpl32 | Rps9 | Tmsb4x |
| Eif1 | Rpl36 | Rps14 | Ubb |
| Fth1 | Rpl37 | Rps15a | mt-Cytb |
| Rpl1 | Rpl37a | Rps19 | mt-Nd1 |
| Rpl11 | Rpl4 | Rps2 | mt-Nd4 |
| Rpl13 | Rpl41 | Rps24 | Rpl26 |
| Rpl14 | Rpl8 | Rps26 | Rps11 |
| Rpl18a | Rplp0 | Rps27 | Rps29 |
| Rpl23 | Rplp1 | Rps27a |  |

**Table S6:** Most enriched GO categories, found by DAVID, for genes whose predicted occurrence is significantly higher than the empirical one. We selected genes with *o*_*i*,predicted_ > *o*_*i*,empirical_ + 0.2 (as highlighted in Figure 7 of the main text). The complete gene lists are available from https://github.com/BioPhys-Turin/Emergent_Laws_in_scRNA-seq_Data/tree/main/MouseCellAtlas/orealopred_lists.

| Organ | Term | Benjamini |
| --- | --- | --- |
| Bladder | extracellular exome | $8.4 * 10^{-17}$ |
| | secreted | $1.3 * 10^{-7}$ |
| | extracellular matrix | $2.3 * 10^{-7}$ |
| | disulfide bond | $1.1 * 10^{-6}$ |
| Brain | blood microparticle | $1.1 * 10^{-5}$ |
| | haptoglobin-hemoglobin complex | $3.1 * 10^{-5}$ |
| | extracellular exosome | $3.6 * 10^{-5}$ |
| | haptoglobin binding | $10^{-5}$ |
| Kidney | extracellular exome | $6.1 * 10^{-12}$ |
| | Antigen processing and presentation | $5.7 * 10^{-5}$ |
| | cytosol | $6.1 * 10^{-5}$ |
| | Chaperone | $2.3 * 10^{-4}$ |
| Liver | Ribosome, cytoplasmic | $10^{-68}$ |
| | extracellular exosome | $1.8 * 10^{-9}$ |
| | haptoglobin-hemoglobin complex | $1.1 * 10^{-4}$ |
| | extracellular space | $1.2 * 10^{-4}$ |
| Muscle | extracellular space | $2.7 * 10^{-10}$ |
| | Ribosomal protein | $4.5 * 10^{-10}$ |
| | Acetylation | $7.6 * 10^{-12}$ |
| | Muscle protein | $4.5 * 10^{-10}$ |
| Pancreas | extracellular space | $6.3 * 10^{-19}$ |
| | Secreted | $2.0 * 10^{-17}$ |
| | Disulfite bond | $2.9 * 10^{-17}$ |
| | signal peptide | $2.2 * 10^{-16}$ |

**Table S7:** Genes having predicted occurrence significantly higher than the empirical one in the Muscle + Brain joint data set. We selected genes with *o*_*i*,predicted_ > *o*_*i*,empirical_ + 0.2.

|  |  |  |
| --- | --- | --- |
| Acta1 | Fos | Plp1 |
| Aplp1 | Ftl1 | Plp1 |
| Apod | Gpx1 | Ptgds |
| Apoe | Hba-a1 | Qdpr |
| Atp1a2 | Hba-a2 | Retnla |
| C1qa | Hbb-bs | Retnlg |
| C1qb | Hbb-bt | S100a8 |
| C1qc | Hexb | S100a9 |
| Camp | Igkc | Scd2 |
| Ccl4 | Lcn2 | Sparc |
| Cd74 | Ltf | Sparcl1 |
| Chil3 | Ly6c2 | Stmn4 |
| Cldn11 | Lyz2 | Tnnc2 |
| Cnp | Mag | Trf |
| Cryab | Mbp | Ttr |
| Cst3 | Meg3 | Tyrobp |
| Ctss | Mmp8 | Wfdc21 |
| Elane | Mobp | Zfp36 |
| Ermn | Ngp | Zfp36 |

**Table S8:**
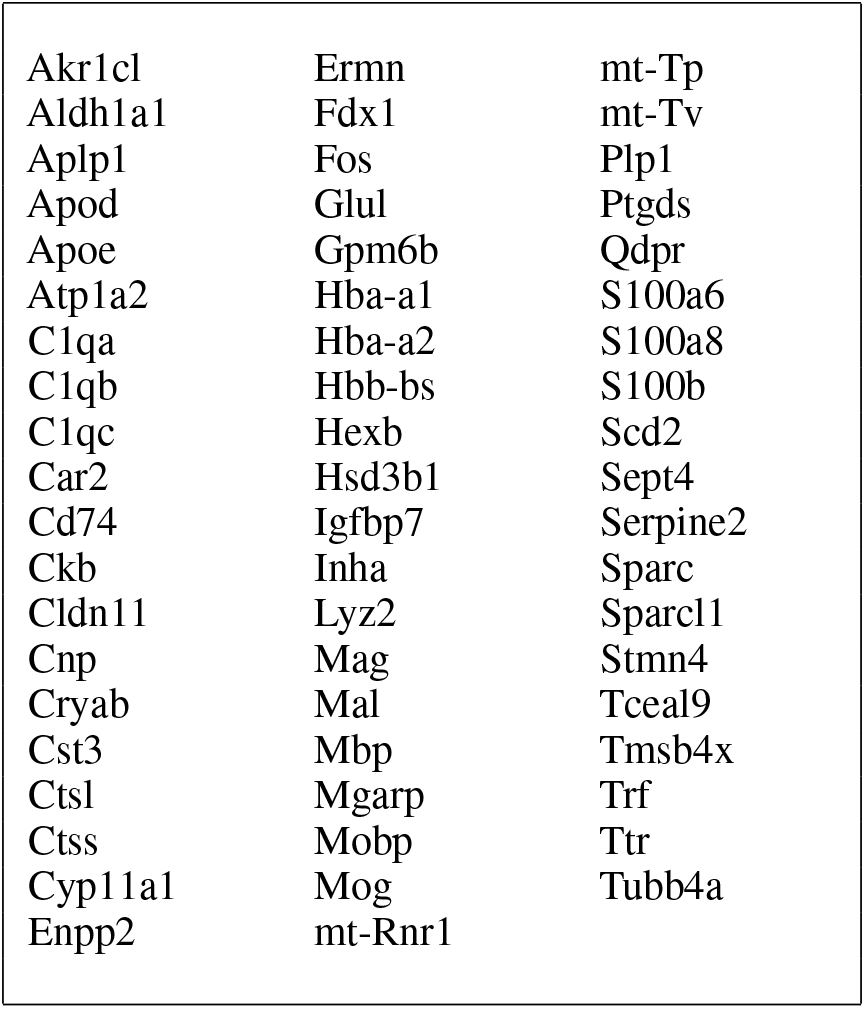
Genes having predicted occurrence significantly higher than the empirical one in the Ovary + Brain joint data set. We selected genes with *o*_*i*,predicted_ > *o*_*i*,empirical_ + 0.2.

**Table S9:** Genes having predicted occurrence significantly higher than the empirical one in the Muscle + Blood joint data set. We selected genes with *o*_*i*,predicted_ > *o*_*i*,empirical_ + 0.2.

|  |  |  |
| --- | --- | --- |
| Anxa1 | H2-Ab1 | Il1b |
| Camp | H2-Eb1 | Lcn2 |
| Ccl5 | Hba-a1 | Ltf |
| Ccl6 | Hba-a2 | Ly6c2 |
| Cd74 | Hbb-bs | Mmp8 |
| Cd79a | Hbb-bt | Mpo |
| Chil3 | Ifitm3 | Ngp |
| Crip1 | Ifitm6 | Plac8 |
| Elane | Igha | Retnlg |
| Gzma | Ighm | S100a6 |
| H2-Aa | Igkc | Wfdc21 |

**Table S10:**
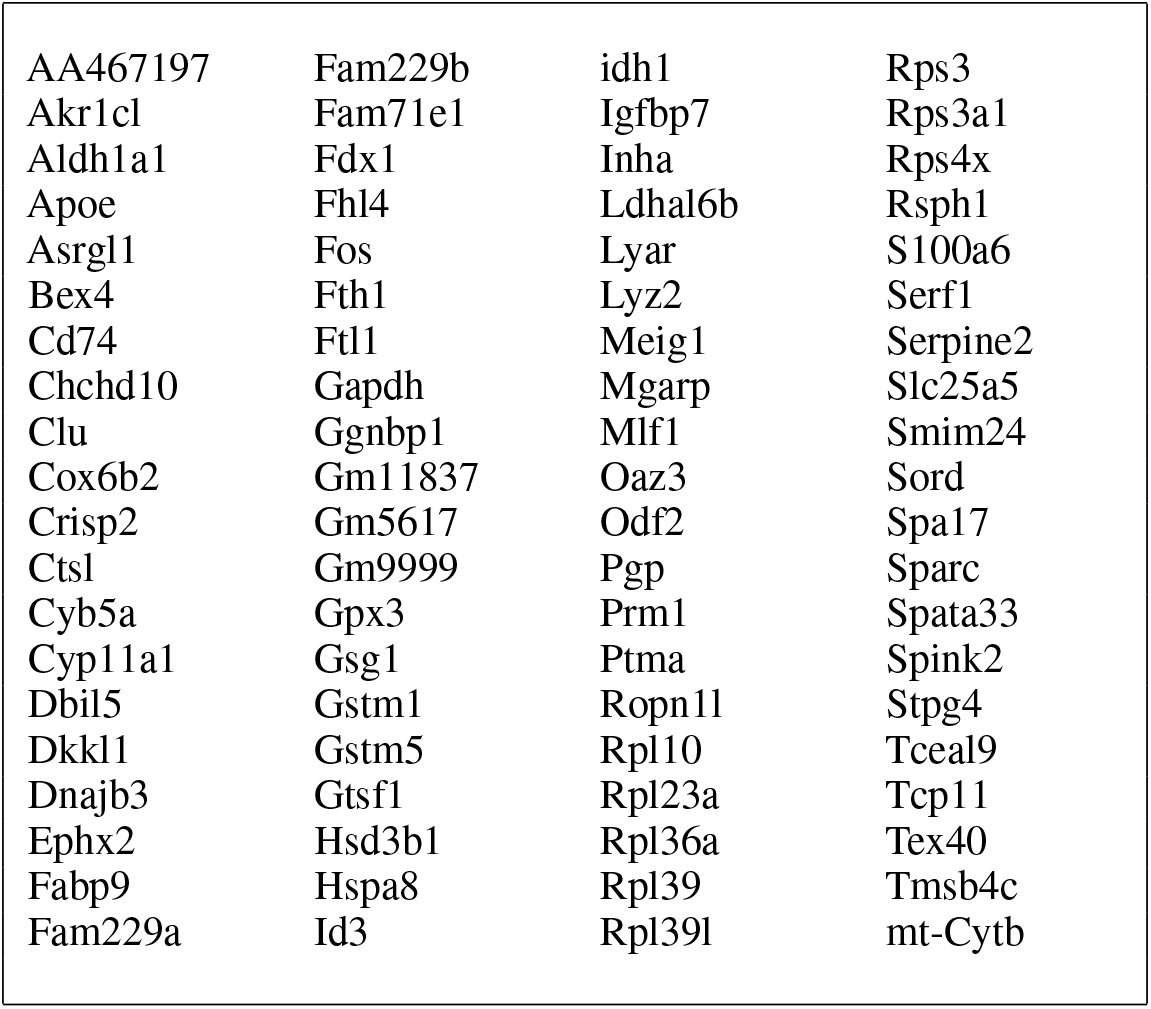
Genes having predicted occurrence significantly higher than the empirical one in the Ovary + Testis joint data set. We selected genes with *o*_*i*,predicted_ > *o*_*i*,empirical_ + 0.2.

**Table S11:** Most enriched categories, found by DAVID, for genes whose predicted occurrence is significantly higher than the empirical one in fibroblasts data set. We selected genes with *o*_*i*,predicted_ > *o*_*i*,empirical_ + 0.4. The gene list is available at https://github.com/BioPhys-Turin/Emergent_Laws_in_scRNA-seq_Data/tree/main/SmartSeq3/opredoreal_lists.

| Term | Benjamini |
| --- | --- |
| extracellular region | $6.2 * 10^{-15}$ |
| mitotic nuclear division | $1.1 * 10^{-13}$ |
| cell cycle | $1.1 * 10^{-13}$ |
| cell division | $1.1 * 10^{-13}$ |

